# Spatial transcriptomic data reveals pure cell types via the mosaic hypothesis

**DOI:** 10.1101/2024.08.09.607193

**Authors:** Yiliu Wang, Christof Koch, Uygar Sümbül

## Abstract

Neurons display remarkable diversity in their anatomical, molecular, and physiological properties. Although observed stereotypy in subsets of neurons is a pillar of neuroscience, clustering in high-dimensional feature spaces, such as those defined by single-cell RNA-seq data, is often inconclusive, with cells seemingly occupying continuous, rather than discrete, regions. In the retina, a layered structure, neurons of the same discrete type avoid spatial proximity with each other. While this principle, which is independent of clustering in feature space, has been a gold standard for retinal cell types, its applicability to the cortex has been only sparsely explored. Here, we provide evidence for such a mosaic hypothesis by developing a statistical point process analysis framework for spatial transcriptomic data. We demonstrate spatial avoidance across many excitatory and inhibitory neuronal types. Spatial avoidance disappears when cell types are merged, potentially offering a gold standard metric for evaluating the purity of putative cell types.

**Significance statement:** While morphologically or molecularly-defined cell types and their taxonomy are a pillar of modern neuroscience, the extent to which their discrete treatment reflects biology is far from settled. A functional hypothesis concerning cortical neuronal cell types with roots in retina research suggests an anatomical test for the existence and identification of pure, discrete cell types. We describe evidence for this decades-old hypothesis in the mouse neocortex and detail the associated computational methodology based on spatial point processes.

## Introduction

Anatomical stereotypy can facilitate the formation of canonical circuits. In the mammalian retina, a part of the central nervous system with a laminar structure like neocortex, different patches of the visual field implement the same transformations to achieve visual equivariance. Hence, neurons of a type should be roughly equally distributed across any central patch of the retina. This requires cells of the same type to *tile* the retinal surface by avoiding being too close to each other. This insight was confirmed by seminal studies,^1, 2, 3^ and now forms a gold standard method of defining pure retinal cell types.^4^ To the extent that neocortical canonical circuits (*i.e*., connectivity motifs repeating across the neocortex) exist, the same argument requires that cells of the same type should be approximately evenly distributed across the cortical sheet. Indeed, Francis Crick used this principle^5^ to hypothesize the existence of on the order of 1,000 cell types in each cortical area (see also^6^).

*Tiling* refers to the approximately non-overlapping nature of the convex hulls of the dendritic arbors of neurons of the same type, like tiles on a bathroom floor. *Mosaic* refers to overlap of dendritic arbors but with cell bodies of the same type avoiding coming too close to each other. In the retina, neurons closer to the photoreceptors tile the surface whereas ganglion cells form a “mosaic”.^4^ Thus, the coverage factor of a type, the average number of neurons innervating a patch, is close to one at the level of the photoreceptors but increases deeper into the retinal circuit. When cell bodies are represented by points in space, both tiling and mosaic patterns bestow upon the resulting point set an organization different from complete spatial randomness (CSR) (Figure 1).

**Figure 1.**
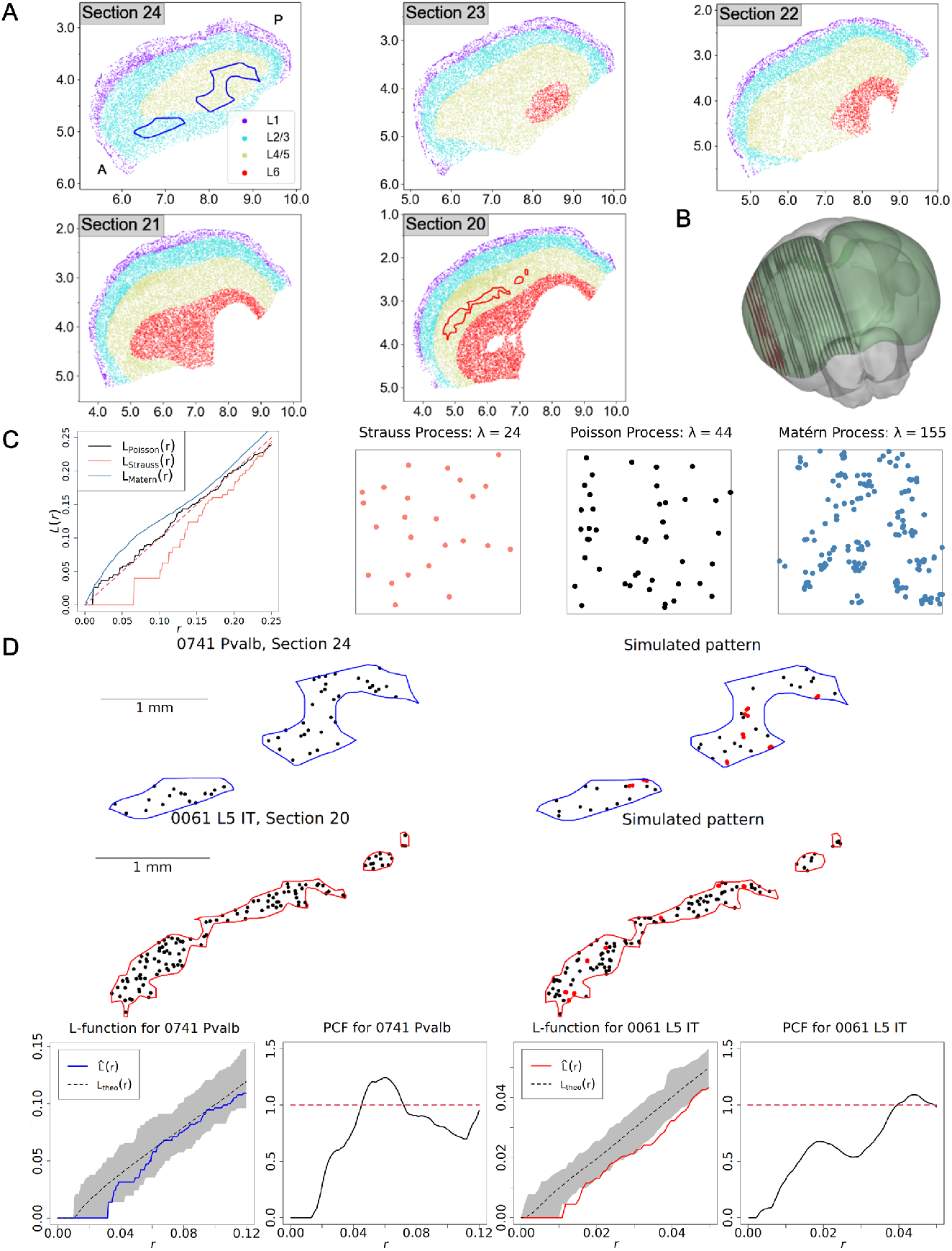
Point process analysis of spatial transcriptomic data with cell type annotations reveals spatial layout of neuronal cell types. **A**. Annotated, detected cells in lateral sections of the mouse brain used in this study, obtained from.^25^ Each dot corresponds to a cell body and the depicted cortical layer annotations suggest the extent of the relevant surfaces for 2D point process analysis. Blue outline: 0741 Pvalb. Red outline: 0061 L5 IT. Top left: brain section number from.^25^ A: anterior, P: posterior. **B**. 3D visualization of locations of the brain sections shown in (A) (red) and those not used (grey) within the CCF atlas.^26, 27^ **C**. Illustration of the L-function for three different point processes with repulsion (*L*_Strauss_ in red), spatial randomness (*L*_Poisson_ in black), and clustering (*L*_Matérn_ in blue). **D**. Illustration of the hypothesis testing procedure for one GABAergic type (0741 Pvalb, *n* = 50) and one glutamatergic type (0061 L5 IT, *n* = 120). Point pairs in the simulated windows that are closer to each other than the corresponding nearest neighbor distance in data are marked in red. Bottom: L- and pairwise correlation functions of the observed patterns (*L*_obs_ in color) together with L-function statistics for the corresponding 199 simulations (Boundaries of shaded region show minimum and maximum L-function values of the simulations.) More simulated patterns are shown in Supplementary Figure S1. All coordinates are in mm. See Table 1 for n-values.

Retinal studies demonstrate that cell death, tangential migration, and dendritic avoidance contribute to the patterning of cell bodies spatial constellation.^7^ While self-avoidance mechanisms regulate neuronal morphology by enabling recognition of sister branches of the same neuron via contact-dependent molecular recognition,^8,9,10,11^ homotypic avoidance mechanisms, such as cell-cell repulsion, regulate the positioning of cells of the same type.^10^ These mechanisms can produce 3-dimensional signatures in the cortex even when the primary effect concerns 2-dimensional positioning in a layer because cortical neurons, unlike retinal ones, do not stratify to occupy thin layers within the cortex.

Astrocytes tile neocortical tissue by forming exclusive territories following migration,^12,13,14,15^ but less is known about the spatial organization of neurons. Maruoka *et al*.^16^ showed microcolumnar organization of layer 5 intra- and extra-telencephalic neurons, demonstrating tangential repulsion of cells in these subclasses. The opposite effect, spatial clustering instead of avoidance, was observed in parvalbumin and somatostatin-expressing neurons.^17,18^ Importantly, these studies concern broad subclasses and not their subsets forming (pure) cell types – subpopulations whose members demonstrate similar phenotypes across all relevant modalities.^19,4,20^ Close approximations to this idealized definition exist in the retina^21^ and the cortex.^22^

Single-cell transcriptomic technology enables the definition of many, putative pure types within those broad subclasses; yet their spatial organization, or lack thereof (*i.e*., spatial randomness), remains unknown. A confounding factor is that clustering in high-dimensional feature spaces is error-prone due to the “curse of dimensionality,” sampling bias, and complicated noise processes, in addition to the ill-posed nature of the clustering objective.^23^ Here, we study a recent whole-brain spatial transcriptomics dataset where over 1,100 genes are profiled in space and a transcriptomic identity (based on reference clustering of single-cell RNA sequencing data^24^) as well as a location within the common coordinate framework (CCF) are assigned to approximately 10 million cells in the mouse^25^ (Figure 1). While this represents a monumental achievement, neither the initial clustering nor the identity assignment steps are immune to the problems listed above. Moreover, 2D (instead of 3D) imaging of brain sections further complicates the analysis of spatial distribution.

We develop a stochastic point process analysis framework to study apparent pairwise short-range (e.g., nearest neighbor) interactions between neurons. Focusing on cell types whose local organization can be studied with statistical rigor, we reveal homotypic avoidance of cell bodies within pure cell types and its loss in mixtures of cell types. We confirm these results in a second dataset with a perpendicular sectioning axis. Our findings support a long-standing hypothesis on the organization of cortical circuitry and outlines a general method to evaluate and validate the purity of putative cell types.

## Results

### Selecting cell types for point process analysis

Three main sources of noise impede point process analysis of spatial transcriptomic datasets. First, reference cell type labels may not be accurate (“label noise”), the root cause of confusion in cell types research. This can be due to inaccuracies in initial single-cell RNA sequencing (scRNA-seq) based clustering,^24^ incorrect mapping of spatial transcriptomic observations to those clusters,^25^ or noise in image-based assignment of gene expression to cells. Second, imperfections in segmentation of cells and profiling their genes can lead to a decrease in the number of annotated cells, acting as another noise source that decreases the apparent intensities of the underlying point processes. We formally prove that label noise and thinning of the annotations bring the underlying process closer to spatial randomness (Theorem 1, Methods). Third, a single 2D projection image is obtained from a slice of tissue that is 10*µm* thick. Thus, the distance between a pair of neurons may appear arbitrarily small in projection images if they are separated mostly along the slicing axis.

To account for these noise sources, we implement two filtering steps (Supplementary Note B, Supplementary Figure S2, Supplementary Table S1). First, we calculate the relative abundance of high confidence cells for each type, defined as cells whose reference mapping confidence exceeds a threshold,^25, 24^ and exclude cell types with low confidence ratios (Supplementary Table S2). This filters against label noise (“label noise filter”).

Our second filter is motivated by the fact that a test may fail to detect homotypic avoidance (*i.e*., reject the null hypothesis of spatial randomness) if the process has low intensity. Concretely, if only a few points exist in a large area, random placement creates patterns, with high probability, without any points close to each other (Supplementary Note C). To understand the effect of the prevalence of a cell type on the ability to reject the null hypothesis, we simulated 2D point processes with different densities (Supplementary Figure S3). We infer the intensity threshold below which the power of our chosen statistical test is insufficient to identify homotypic avoidance (Methods, Supplementary Figures S4, S5). For each type, we only consider brain sections with sufficient number and intensity (average density of cells) of high confidence cells to act as a filter against low intensity patterns (“minimal intensity filter”).

Lastly, instead of filtering out cell types with dense point patterns that are prone to distortions due to 3D to 2D projection (e.g., excitatory populations), we account for the impact of the projection via the setup of the statistical test as we introduce next.

### Statistical testing for spatial organization of cell bodies

To analyze the local organization in the point pattern of cell bodies in 2D brain sections, we first identify a set of regions tightly covering the points belonging to the transcriptionally defined cell type of interest, to avoid falsely assigning significance to global patterns of cell abundance (Supplementary Note I, Supplementary Figures S6, S7, S8, S9, and S10). We calculate the average normalized abundance of points from a given point across these regions as a function of distance from this point, known as the L-function^28^ (Figure 1C,D).

For each cell type, we simulate the null hypothesis of CSR with the same number of points multiple times in the same region. Noting that the L-function, expressing normalized intensity as a function of distance (Methods), of a Poisson process concentrates around the identity function (i.e., *L*(*x*) = *x* as in Figure 1C), we use the Diggle-Cressie-Loosmore-Ford (DCLF) test^28, 29^ to measure the significance of one-sided deviations from CSR based on the area between the measured and simulated L-functions. While a deviation below the *L*(*x*) = *x* line at *x*_0_ implies avoidance of neighbors up to a distance *x*_0_, a deviation above it implies spatial clustering (Figure 1C).

CSR can be trivially rejected given that cell bodies are non-penetrating bodies of finite size so that points cannot be arbitrarily close to each other in 3D. We address this by applying the DCLF test only across a range of interactions that excludes this shortest scale that we term the *hardcore distance*. We obtain an unbiased distribution of soma sizes from a volume electron microscopy dataset^30^(Supplementary Note A, Supplementary Figure S11). Moreover, we show that the signature of the hardcore separation between GABAergic cells can extend beyond the hardcore distance, but the projection onto two dimensions effectively destroys the hardcore separation for glutamatergic subpopulations (Supplementary Figure S12), which are more numerous (Figure 1D). Therefore, for GABAergic types, we generate the benchmark 2D simulations from a process with the finite minimal distance, known as the Strauss hardcore process, instead of a canonical Poisson point process (for Glutamatergic types, see Methods and Supplementary Table S3).

We take advantage of the fact that multiple brain sections can pass the noise filtering steps for a given cell type by applying a meta analysis of the DCLF test results using Fisher’s method^31^ (Supplementary Note D). Thus, we formally test the null hypothesis that the point pattern formed by cells display spatial randomness in *all* eligible brain sections (Figure 2, blue disks).

**Figure 2.**
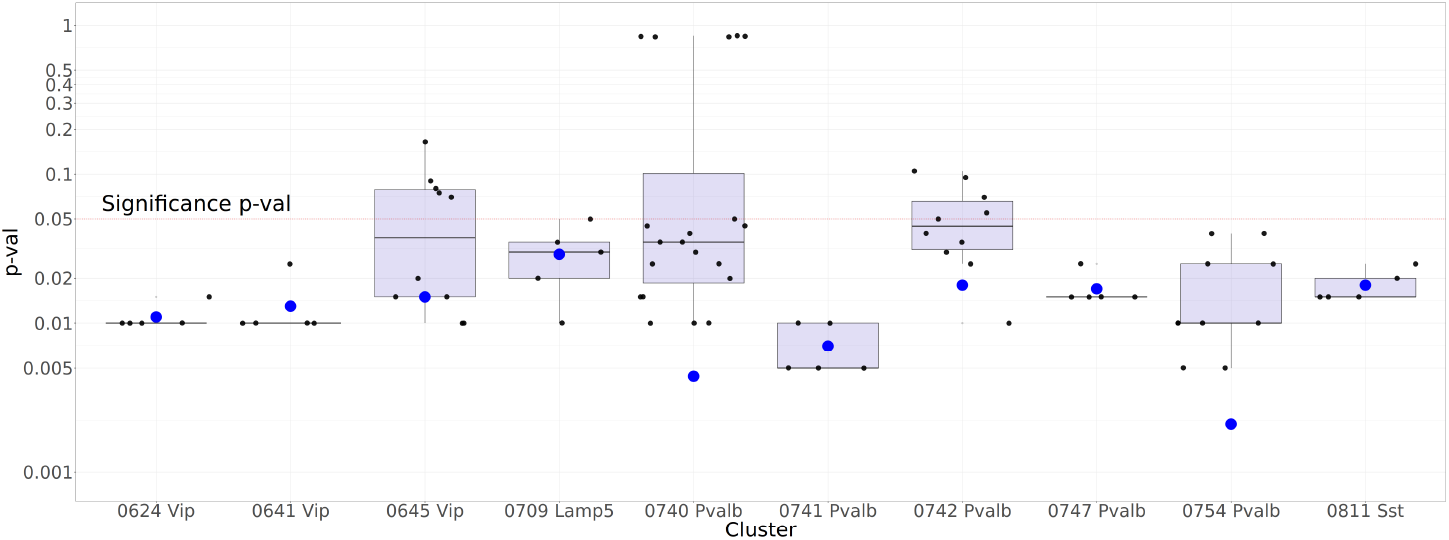
Statistical testing rejects complete spatial randomness (CSR) for all ten GABAergic types passing the filtering steps, at a significance level of 0.05 (1-sided DCLF test). Small, black dots indicate p-values over individual sections for the transcriptomic types. The test is repeated over multiple brain sections using 5 Monte-Carlo simulations with different seeds, each with 199 point pattern samples. Large, blue dots indicate the aggregated p-values calculated according to Fisher’s method. Whisker plot indicates median (black line) as well as first and third quartiles (blue shaded region). When multiple sections are admitted for a cell type, Fisher’s method tests for the weaker hypothesis that points (somata) are distributed according to CSR in all sections. All aggregated p-values remain significant (*i.e*., < 0.05) following correction for multiple testing at a false discovery rate of 0.05, using the Benjamini-Hochberg procedure (Supplementary Tables S4 and S5). See Table 1 for n-values.

### Cells of the same transcriptomic type display homotypic avoidance in neocortex

Our selection procedure picks 28 neuronal cell types across 5 (out of 23; Figure 1B) sagittal brain sections for formal statistical testing. Table 1 displays the results of the DCLF test on these cell types, corrected for multiple testing by the Benjamini-Hochberg procedure.^32^ This controls the expected proportion of false discoveries within all tested cell types, different from Fisher’s method that performs a meta-analysis over the different sections. (See Supplementary Note H and Supplementary Table S6 for a list of differentially expressed genes for each studied cell type.)

**Table 1.** A majority of Glutamatergic clusters and all GABAergic clusters display homotypic avoidance. Sixteen out of 18 excitatory and all 10 inhibitory cell types display statistically significant homotypic avoidance. Our method infers that the two greyed-out excitatory types with low statistical significance are either mixtures, each containing more than one true cell type, or due to over-splitting of a genuine cell type. p-values across different brain sections are combined by Fisher’s method and corrected for multiple testing. `‘p-value’` denotes the fraction of CSR simulations that display as or more extreme homotypic avoidance according to the one-sided DCLF test. `‘Area’` denotes the total hull area of the underlying point pattern. `‘R’` is the approximate interaction range that serves as the upper value for the DCLF test. `‘%’`: the average percentile of nearest neighbor distances *<* 10*µm*, ‘*λ*’: the average intensity, *i.e*., cell density, across qualified sections, `‘conf’`: the average classification confidence based on the reference study,^25^ `‘n’`: the total number of cell bodies. See Supplementary Tables S4 and S5 for the p-values for individual cluster-section combinations, Supplementary Note F and Supplementary Figures S13 and S14 for the pairwise correlation functions of individual clusters, and Supplementary Table S6 for the list of differentially expressed genes in each studied cell type according to Ref.^24^

| Type | Section | p-value | Area(mm<sup>2</sup>) | R(μm) | % | $\lambda$(1/mm<sup>2</sup>) | conf | n |
| --- | --- | --- | --- | --- | --- | --- | --- | --- |
| 0109 L2/3 IT CTX Glut_2 | 20–24 | 7.5e-08 | 4.04 | 18 | 8.53 | 1053 | 0.88 | 4261 |
| 0074 L4/5 IT CTX Glut_1 | 20–24 | 1.1e-06 | 1.52 | 27 | 3.96 | 476 | 0.75 | 649 |
| 0079 L4/5 IT CTX Glut_2 | 20,21,24 | 4.1e-05 | 1.27 | 16 | 11.68 | 1060 | 0.74 | 1404 |
| 0455 L6 CT CTX Glut_5 | 20,21 | 3.9e-04 | 0.61 | 24 | 2.68 | 506 | 0.77 | 332 |
| 0116 L2/3 IT CTX Glut_3 | 21,23,24 | 1.1e-03 | 0.30 | 20 | 9.59 | 816 | 0.78 | 268 |
| 0092 L4/5 IT CTX Glut_4 | 23,24 | 3.0e-03 | 0.38 | 50 | 0.00 | 243 | 0.71 | 88 |
| 0061 L5 IT CTX Glut_3 | 20 | 5.0e-03 | 0.44 | 38 | 0.00 | 271 | 0.78 | 120 |
| 0114 L2/3 IT CTX Glut_3 | 24 | 5.0e-03 | 0.10 | 29 | 3.08 | 643 | 0.79 | 65 |
| 0443 L6 CT CTX Glut_2 | 21 | 5.0e-03 | 0.31 | 29 | 1.40 | 456 | 0.77 | 143 |
| 0086 L4/5 IT CTX Glut_3 | 23,24 | 6.6e-03 | 0.48 | 53 | 0.00 | 112 | 0.84 | 44 |
| 0375 L5 ET CTX Glut_5 | 23 | 0.011 | 0.44 | 29 | 2.70 | 340 | 0.80 | 148 |
| 0026 L5/6 IT TPE-ENT Glut_2 | 23 | 0.015 | 0.44 | 49 | 0.00 | 133 | 0.70 | 59 |
| 0439 L6 CT CTX Glut_1 | 21 | 0.024 | 0.36 | 32 | 9.52 | 347 | 0.75 | 126 |
| 0007 IT EP-CLA Glut_1 | 20 | 0.027 | 0.10 | 28 | 3.77 | 518 | 0.90 | 53 |
| 0359 L5 ET CTX Glut_2 | 23 | 0.039 | 0.10 | 41 | 0.00 | 165 | 0.83 | 16 |
| 0003 CLA-EPd-CTX Car3 Glut_1 | 20,21 | 0.044 | 1.05 | 40 | 9.25 | 198 | 0.78 | 207 |
| 0453 L6 CT CTX Glut_5 | 20 | 0.061 | 0.05 | 22 | 0.00 | 275 | 0.80 | 15 |
| 0004 CLA-EPd-CTX Car3 Glut_1 | 23, 21 | 0.14 | 1.17 | 51 | 3.75 | 103 | 0.84 | 122 |
| 0754 Pvalb Gaba_8 | 20, 21 | 0.002 | 2.58 | 92 | 1.67 | 43 | 0.86 | 100 |
| 0740 Pvalb Gaba_3 | 20,21,23,24 | 0.004 | 1.85 | 59 | 2.94 | 84 | 0.81 | 158 |
| 0741 Pvalb Gaba_3 | 24 | 0.007 | 0.76 | 68 | 0.00 | 66 | 0.84 | 50 |
| 0624 Vip Gaba_1 | 24 | 0.011 | 0.94 | 120 | 0.00 | 12 | 0.86 | 11 |
| 0641 Vip Gaba_5 | 22 | 0.013 | 0.23 | 120 | 0.00 | 17 | 0.72 | 4 |
| 0645 Vip Gaba_6 | 20,24 | 0.015 | 1.79 | 120 | 0.00 | 14 | 0.82 | 26 |
| 0747 Pvalb Gaba_4 | 20 | 0.017 | 0.76 | 120 | 0.00 | 20 | 0.82 | 15 |
| 0742 Pvalb Gaba_3 | 23,24 | 0.018 | 1.14 | 93 | 0.00 | 39 | 0.89 | 44 |
| 0811 Sst Gaba_13 | 21 | 0.018 | 1.06 | 120 | 0.00 | 7 | 0.74 | 7 |
| 0709 Lamp5 Gaba_1 | 22 | 0.029 | 0.28 | 82 | 0.00 | 40 | 0.76 | 11 |

To study the noise processes introduced earlier, we also calculate three statistics for each cell type. (i) The average percentile of nearest neighbor distances that are *<* 10*µm* relates to the severity of distortions due to observing a 3D brain section projecting onto a 2D plane. (ii) The average intensity across qualified sections is informative concerning cell types for which the DCLF test may fail to detect homotypic avoidance due to either a low coverage factor or missing cells. (iii) Lastly, the classification confidence refers to the average confidence score of that cell type label from the original study.^25^ A lower average classification confidence implies that the cluster may suffer more from label noise. We also report the sections, the total hull area and the total number of points (summed over qualified sections) of the underlying point pattern, and the upper limit of the interaction range we use to conduct the DCLF test. Homotypic avoidance behavior, when it exists, should be observable over short distances for pairwise interactions. Therefore, we define the upper limit as the median nearest neighborhood distance across sections where the cluster passes the noise and minimum intensity filters.

### Glutamatergic clusters

18 excitatory neuronal clusters pass our filtering steps across 40 sections (Table 1). Of these, the null hypothesis of CSR is rejected in 16 clusters, in favor of homotypic avoidance with *p* = 0.05. Note that the two clusters that fail to reach this level of statistical significance (0453 L6 CT CTX Glut 5 and 0004 CLA-EPd-CTX Car3 Glut 1) have relatively low intensities in their selected regions. They may also correspond to a mixture of pure types, either due to under-splitting or mistakes in cluster boundaries in the reference study.

From a geometric perspective, ∼11,500 non-overlapping disks with ∼10*µm* diameter can be fitted into a 1*mm*^2^ square.^33^ Therefore, when somata are regularly spaced, the intensities listed in Table 1 – peaking at 1060 cells/mm^2^ – suggest that the homotypic avoidance interaction ranges are greater than the hardcore separation due to the soma size. That is, the interaction ranges of all clusters for which CSR is rejected are greater than the hardcore separation due to non-penetrating cell bodies.

### GABAergic clusters

Table 1 includes the results of the same analysis for the 10 GABAergic clusters that pass the noise filtering step across 20 sections. Complete spatial randomness (CSR) can be rejected for all 10 (at *p* = 0.05): indeed, they all show homotypic avoidance (Figure 2).

Since there are fewer GABAergic than Glutamatergic cells, the noise introduced by 2d projection imaging of the 10*µm*-thick brain sections, which can distort the interpoint distances, is less of a concern for point process analysis of GABAergic clusters (Table 1). On the other hand, this makes them more vulnerable to thinning of the population (e.g., due to exclusion of cell annotations failing quality control). As discussed above, the power of the statistical test in identifying patterns diminishes with decreasing intensity.

### A comparative perspective via the density recovery profile

Studies in the retina use the density recovery profile (DRP), which is an intuitive histogram function closely related to the pairwise correlation function, to assess homotypic avoidance of subpopulations.^34,3,21,35^ While we base our analysis on the L-function, which is superior to the DRP for hypothesis testing (Methods, Ref.^28^), we also seek to obtain a comparative perspective by computing DRPs of example cortical cell types. We observe that the DRPs of cortical populations are qualitatively similar to those of retinal populations (Supplementary Note G, Supplementary Figure S15). This similarity extends to populations that are mixtures of cell types, for which the DRP does not display an initial low density region. Indeed, Supplementary Figure S15 is comparable to Figure 2 of Ref.^3^ as a whole.

### Spatial clustering of GABAergic neurons at the subclass level

Given that many cell types defined at the finest transcriptional level avoid each other (Table 1), we next study whether this is true at coarser levels of the transcriptional hierarchy.^24^ Indeed, GABAergic neurons of the same subclass (e.g., all Pvalb neurons) show the opposite pattern, clustering together, possibly to improve the impact of their inhibitory function.^17, 18^

We formally test spatial clustering for the Vip, Lamp5, Pvalb, and Sst subclasses. For each, we consider sections where at least one cell type belonging to the subclass passes the intensity filter (we exclude the 0641 Vip type in Section 22 from this analysis because it has too few samples (four) to impact the mixture statistics). We use the 1-sided DCLF test for spatial clustering (e.g., *L*(*x*) *> x*) at the subclass level with the same benchmark process for spatial randomness as before, in the range between the hardcore separation distance and 20*µm*.^17^

Consistent with previous findings,^17,18^ all inhibitory subclasses cluster at near distances (ca. 20*µm*) with high statistical significance, and are distributed randomly for larger distances (Supplementary Tables S7, S8 and Figure S16).

### Segregation of glutamatergic neurons at the subclass level

While GABAergic neurons cluster at the subclass level, this is not the case for excitatory, glutamatergic cells. Figures 3A and 3B show the cell distribution of L2/3 IT neurons and Pvalb neurons in sagittal section 24, highlighting the dominant clusters for both excitatory and inhibitory cells that are non-randomly distributed (p=0.05). Glutamatergic types do not mix as well as GABAergic types. We quantify this observation by defining a segregation index for each subclass as the unnormalized *k*-nearest-neighbor correlation,^36^ which is the mean fraction of cells among the *k*-nearest neighbours (*k*=5) in the same subclass having the same cell type label as the reference cell acorss sections. Hence, an index close to 1 indicates a well-separated subclass while an index close to 0 indicates complete spatial mixing. Cells of the three L2/3 IT types (Figure 3B) occupy different regions at the subclass level (Mean segregation index: 0.77), while spatial segregation is much weaker for Pvalb clusters (Mean segregation index: 0.40). Indeed, in general, GABAergic cell types mix spatially with each other while glutamatergic cell types are typically more segregated *within the context of their subclass* (Figure 3C). Thus, the anatomical location of the cluster does not offer a way to verify fine-scale GABAergic types.

**Figure 3.**
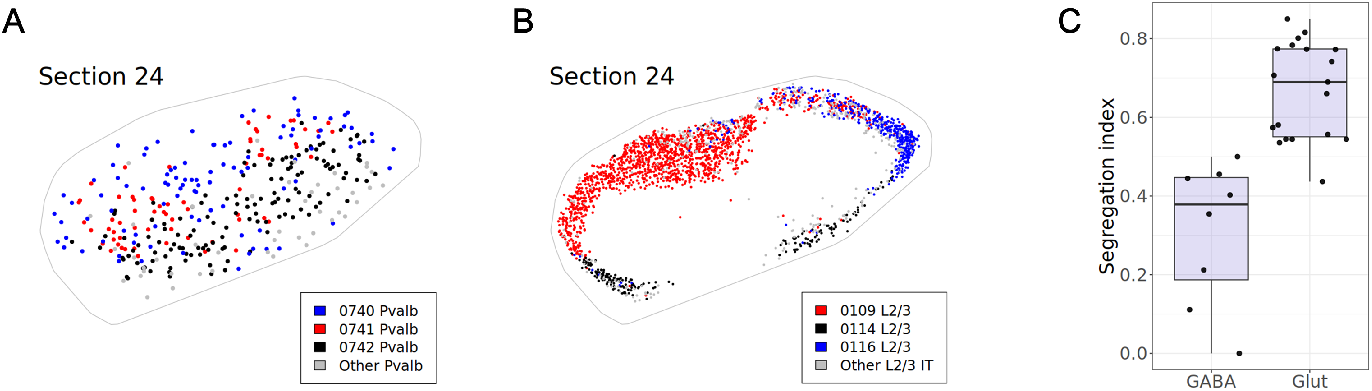
Segregation effect at the subclass level. **A**. Spatial distribution of Pvalb types that co-mingle. **B**. Spatial distribution of L2/3 IT types that predominate in separate regions. **C**. Segregation indices organized by neuronal classes for the five sagittal brain sections 20-24. Each dot indicates the index corresponding to one subclass in one section, calculated for sections where at least one cell type belonging to the subclass rejects CSR.

### Homotypic avoidance disappears in mixtures of transcriptomically similar cell types

We formally test homotypic avoidance at the subclass level for the four GABAergic subclasses. We conduct the DCLF test between the hardcore separation distance and the upper interaction range derived for each subclass and find that none of the subclasses display statistically significant homotypic avoidance for any of the five sections (Supplementary Tables S7 and S8).

This result puts forth the interesting possibility that, as in the retina, homotypic avoidance disappears as one moves up the transcriptomic hierarchy, offering a gold standard test of what defines a cortical cell type, sidestepping some of the difficulties associated with clustering in high-dimensional feature spaces. To elucidate the transition from statistically significant homotypic avoidance at the level of transcriptomic types to its lack at the subclass level, we merge groups along the transcriptomic hierarchy below the subclass level, referred to as supertypes.

A confounding factor is that many types are not spatially intermixed at the supertype level (e.g., 0741 Pvalb and 0742 Pvalb in Figure 3). An exception is 0740 Pvalb and 0742 Pvalb, which strongly co-exist in two regions (SSs4 and VISC5) in section 23 (Supplementary Note E, Supplementary Table S9): indeed, CSR is rejected for both clusters at p=0.05 (Table S5); yet merging them into a single group destroys homotypic avoidance, supporting the hypothesis that they constitute genuine cell types (Supplementary Figure S17b).

We expand this analysis to all supertypes and subclasses in the filtered dataset. For example, Pvalb 0740, 0741 and 0742 all belong to the same Pvalb supertype, 0207. Yet CSR is rejected (at a Fisher-aggregated p-value of 0.13) when applying the same region selection method and hypothesis test to 0207. To facilitate the interpretation, we define the concept of effective cluster count (ECC) as the exponential of the entropy of the mixture pattern, which calculates an information-theoretic estimate for the number of clusters present in the sample pattern (Methods). For a mixture consisting of samples from *N* clusters, the ECC value has a minimum of 1, denoting the mixture has samples from a single cluster, and a maximum of *N*, denoting all clusters are equally represented in the mixture. The higher the ECC value, the more mixed the sample population is (Figure 4A). For instance, the supertype 0207 Pvalb has an ECC value of 2.84, implying that it is dominated by roughly three clusters.

**Figure 4.**
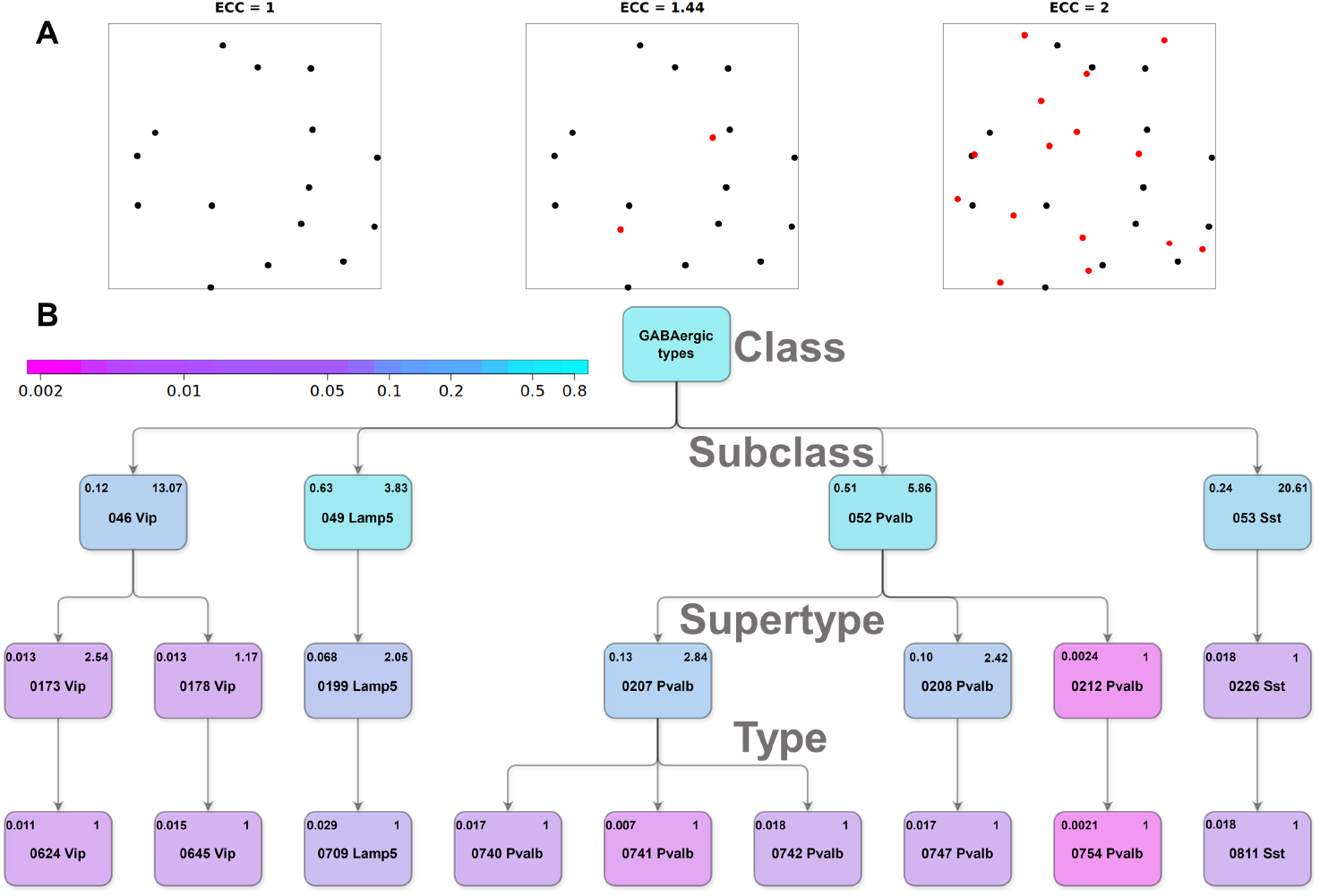
Increase of p-values in mixture types. **A**. *Effective cluster count* (ECC) values for N=2 point patterns. ECC has a minimum of 1 when there is a single type and a maximum of *N* when *N* types are spatially intermixed. The larger ECC, the more mixed the underlying population. **B**. Tree hierarchy diagram for GABAergic types. Nodes are color colored according to the p-values of the CSR test, with their numerical values in the upper left and their ECC values in the upper right corner. When moving up the hierarchy, pure cell types merge into supertypes and subclasses, and spatial avoidance disappears. The transition is governed by the ECC value of the population.

Figure 4 shows the statistical results and ECC values for all GABAergic supertypes and subclasses that reject CSR (1st level: class, 2nd level: subclasses, 3rd level: supertypes, leaf level: types). Some supertypes have ECC around 1 (0178 Vip, 0212 Pvalb and 0226 Sst), implying domination by one cluster in the selected region. On the other hand, all subclasses have ECC values much greater than 1. While we use all cells belonging to a group for calculating ECC and p-values, individual cell types within a group may be excluded by the filtering steps (due to sample scarcity or unconfident cell type assignment). As an example, the supertype 0208 Pvalb has an ECC value of 2.42, but only one of its cell types (0747) are shown in Figure 4B. For this supertype, the cell counts in Section 20 within the supertype’s observation window are given as *n*_0747_ : *n*_0743_ : *n*_0746_ = 19 : 9 : 3. In contrast, the cell counts in that same section for the supertype 0207 Pvalb are *n*_0740_ : *n*_0741_ : *n*_0742_ = 98 : 39 : 39, so that all of its individual types could be included in Figure 4B.

Importantly, as we go up the tree hierarchy, we see an increase in p-values whenever there is an increase in ECC. This suggests a potential rule expressing the purity of a cell type: the test does not reject CSR for any subclass and rejects it for all transcriptomic types in Figure 4. Moreover, CSR is not rejected for supertypes when the corresponding ECC is 2 or higher. The only exception is the supertype ‘0173 Vip’ (one-sided DCLF p=0.013, ECC=2.54). While noise sources thinning the underlying process can explain this exception where the test is performed on only 11 ‘0173 Vip’ cells, an alternative explanation is that the reference hierarchy oversplits a single genuine cell type (Supplementary Figure S18a and S18b).

### A coronal dataset supports homotypic avoidance

The mouse brain is the longest along the anterior-posterior axis. Moreover, sagittal cuts, as evaluated up to now, follow the cortical sheet more closely in lateral areas compared to coronal cuts in dorsal areas and allow for profiling larger regions to increase the statistical strength of point process analysis based on 2D cuts. Nevertheless, the coronal whole-brain spatial transcriptomic dataset that was also introduced by Ref.^25^ provides an opportunity to test our findings in a different brain region of a different animal. Therefore, we test homotypic avoidance in the visual cortex by studying the most posterior three slices in this coronal dataset (Supplementary Note J, Supplementary Figure S19). By following the same computational pipeline, without tuning any parameters, we arrive at qualitatively identical results (Supp. Table S10): 21 out of 25 Glutamatergic cell types that pass the filtering steps and all 5 GABAergic types that pass the filtering steps display statistically significant homotypic avoidance. Moreover, the inferred interaction ranges and point process intensities are consistent across Tables 1 and S10. The cell types for which homotypic avoidance does not appear statistically significant have relatively fewer samples within the Glutamatergic class. Finally, these cell types are different across the two datasets (L6 types in the sagittal dataset vs. L5 types in the coronal dataset), suggesting the possibility that failure to establish significant homotypic avoidance for those types is due to various noise processes and statistical strength, rather than the underlying biology.

## Discussion

Morphologically defined neuronal cell types formed the basis of Cajal’s exposition of the nervous system.^37^ While a cell-type-centric view gradually fell out of fashion, the last few decades witnessed a forceful return of this concept. Identifying and accessing cell types has received much attention and funding.^38^ Indeed, studying “identical” cells using transcriptional, morphological, developmental, and functional modalities enables operational access and control as well as conceptual leaps in our understanding of nervous system function.^39^

However, this promise can only be realized when the underlying cell type definitions are pure and are neither fragmented nor lumped. In this paper, we provided evidence for the long-standing hypothesis that, like the retina, neocortical neurons of the same type display a mosaic pattern and that this property disappears for mixtures of cell types. This bestows upon the point pattern of cell bodies a distinct statistical signature which can be used to assess the purity of cell types.

Our findings support the existence of cortical columns that implement a canonical circuit capable of a wide range of computations.^40^ This requires the availability of all circuit components (*i.e*., cell types) in each column. Therefore, efficient placement suggests that cells of the same type avoid each other to maintain a relatively uniform coverage. This argument led to the speculation that there may be on the order of 1,000 cell types in cortex,^5^ roughly in agreement with recent transcriptomic studies.^24^ Discovering and verifying these cellular components will be a huge step forward towards revealing such canonical connectivity motifs.^41^ Experience constantly modifies the phenotype, overlaying continuous factors of variability onto discrete identities.^42^ In addition, the “curse of dimensionality” makes it harder to uncover the discrete factors as the feature dimensionality grows to 10,000 or more of expressed genes or RNA transcripts. The statistical procedure put forth here can verify the purity of a proposed cell type without relying on the features used by the clustering algorithm ^43^ as it only depends on the theory of point processes in the plane. That is, if two distinct populations of cells are truly discrete cell types, then they should display repulsion and violate complete spatial randomness (CSR) when treated separately, but not when lumped into a single class, except for clustering at short distances for GABAergic cells.

The original homotypic avoidance hypothesis concerns positioning of cells on a sheet of tissue.^5, 6^ Here, we argue that many homotypic avoidance mechanisms (e.g., diffusion and detection of molecules) should create signatures throughout the sheet as well, especially when the neuronal arbor does not stratify in one thin layer. Hence, we extend the original hypothesis into the third dimension. Nevertheless, if the primary effect concerns positioning in 2D, it is reasonable that the effect size in the third dimension may be smaller. This suggests that larger effect sizes, lower p-values, and more sensitive tests can be obtained when true 3D single-cell molecular datasets become available, offering a strong test of our main findings. 3D imaging will also remove distortions due to 2D imaging of tissue. Alternatives, such as volumetric electron microscopy datasets, offer impeccable spatial characterization, but their field of view remains limited and detailed cell type proposals are not yet available for them.

A source of noise in our study are potential inaccuracies in the underlying transcriptomic cell type definitions or mistakes in inferring those labels from spatial transcriptomic observations with limited gene sets and batch effects. While the reference study associates each putative transcriptomic type with at least one differentially expressed gene^24^ (Supplementary Table S6), noise in high-dimensional expression vectors can create spurious patterns of differential expression. The method presented here will not reject the null hypothesis with a mixture population (*i.e*., under-splitting or incorrect boundaries in the feature space between neighboring cell types) or it may exclude the population from further analysis due to low intensity (that is, low cell count) when the true cell type is over-split into subsets. Note that the number of genes differentiating a cell type from its close peers in Ref.^24^ is smaller than that for retinal bipolar cells,^44^ yet similar to that differentiating retinal ganglion cells,^45^ the only retinal cells projecting outside the local circuitry.

The extent to which the methodology of this study can be fruitfully applied to subcortical regions that have a nuclear structure, such as the hypothalamus, is an open question. On the other hand, the implications of homotypic avoidance are clearer in the case of the cortical sheet that is much more spatially extended than thick. We call on the community to test their own cortical cell types datasets for spatial mosaicism, using our freely available computational pipeline. Ultimate validation of this hypothesis will come from dense volumetric datasets.

## Data and code availability

We study two recent whole-brain spatial transcriptomics datasets by (25). The datasets are publicly available at https://alleninstitute.github.io/abc_atlas_access/. The code for reproducing all figures in the main text and supplementary material is available at https://github.com/yiliuw/mosaics.

## Acknowledgements

We wish to thank the founder of the *Allen Institute*, P. G. Allen, for his vision, encouragement and support. The original motivation to pursue this question arose from discussions between Francis Crick and C.K. many decades earlier. We wish to thank Zizhen Yao, Michael Kunst, and Kelly Jin for useful discussions, Forrest Collman and Leila Elabbady for guidance on the MICrONS dataset, Forrest Collman and Stephen Smith for valuable feedback on the manuscript. Y.W. is a *Shanahan Foundation Fellow*.

## Methods

### Finite Poisson and Strauss point processes

We consider a finite point pattern ***x*** = (*x*_1_, …, *x*_*n*_) observed in a bounded window *W*. The probability density function of finite point processes are defined with respect to a Poisson process of unit intensity.^28^

#### Poisson process

The homogeneuous Poisson process with intensity *β* has the following probability density,

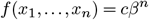

where *c* is a normalizing constant such that *c* = exp ((1 − *β*) |*W*|). Note that the probability density does not depend on the locations of the points, so that different points are independently and uniformly distributed. The conditional Poisson process with *n* points is the binomial process, with probability density *f*_*n*_(*x*_1_, …, *x*_*n*_) = 1*/*|*W*|^*n*^.

#### Matérn process

A Matérn process can be built from the homogeneous Poisson process, capturing the intuition of spatial clustering. A parent process is sampled from a homogeneous Poisson process with intensity *κ*. Then, each parent point has a Poisson(*µ*) number of offspring, independently and uniformly distributed in a disc of radius r centered around the parent. The Matérn process consists of all the offsprings only.

#### Strauss process

The Strauss process has the following probability density function,

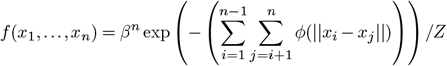

Where

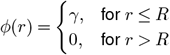

and *Z* is an intractable normalizing constant. The Strauss process is called a hardcore process if *γ* = ∞ for *r* ≤ *R*, implying that no pairs of points that are closer than the distance *R* is allowed. The Strauss process has three important parameters, *β, γ* and *R*. Note that for Strauss processes, *β* is called the self-potential or chemical activity, not the intensity of the model. The parameter *γ* such that *γ* ≥ 0 is the interaction parameter controlling the strength of inhibition and *R* is the range of interaction.

Due to the intractable normalization factor, it is hard to work directly with the density function. Instead, we consider the Papangelou conditional intensity function, defined as *λ*(*u*|***x***) = *f* (***x*** *∪{u}}*)*/f* (***x***).

Let

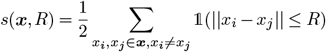

denote the number of pairs of points with an interpoint distance of *R* or less and *t*(*u*, ***x***, *R*) = *s*(***x*** ∪{*u*}, *R*) −*s*(***x***, *R*) denote the number of points with an interpoint distance of *R* or less to a typical point *u*. We can write the Papangelou conditional intensity function of the Strauss process above as

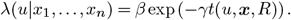

#### Mixture process

We consider a superposition of a Strauss process with interaction strength *γ* and a homogeneous Poisson process. The following theorem shows that the resulting mixture point process can be approximated by another Strauss process with a weaker interaction strength *η < γ*.

##### Theorem 1

Let ***x***_**1**_ be a realization of a Strauss process with interaction strength *γ* ≥ 0, interaction range *R* and self-potential *β* in a study region *W* ∈ ℝ^*d*^. Let ***x***_**0**_ be another point pattern in the same study region. We assume ***x***_**0**_ is a realization of a Poisson process with intensity *aβ* where *a >* 0. The mixture process, defined as *x* = ***x***_**1**_ ∪ ***x***_**0**_ is also a Strauss process with the same self-potential *β* and interaction range *R*, but with a weaker interaction strength *η*.

Moreover, when ***x***_**0**_ and ***x***_**1**_ are finite processes with *n*_0_ and *n*_1_ points respectively and *α* = *n*_0_*/n*_1_ denotes the noise/label ratio,

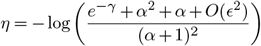

for some small *ϵ >* 0 with *ϵ* = (1 *−e*^*−γ*^)*/g*_1_, *g*_1_ *≥* 0.

We show the proof in Supplementary Note C.

### Summary functions for spatial point processes

Here we provide brief descriptions of common summary functions. We refer to Ref.^28^ for details.

#### The nearest-neighbor function

The nearest-neighbor function *G*(*r*) is defined as the distribution function of the distance from a typical point to its nearest neighbor. It usually serves as the first step in analysis as it is easy to calculate. However, as it only considers the nearest neighbor (short-sighted), it may not be very useful in many cases.

#### Ripley’s K-function

Ripley’s K-function calculates the average number of points found within distance *r* from a typical point in the spatial point process, normalized by the overall intensity of the process.

#### Pairwise correlation function (PCF) and density recovery profile (DRP)

The PCF and the DRP both characterize the pair-distances of a spatial point process and are proportional to the derivative of Ripley’s K-function, *K*(*r*). In the planar case, the PCF is defined as *g*(*r*) = *K*^*′*^(*r*)*/*2*πr* for *r* ≥ 0. For homogeneous Poisson processes, *g*(*r*) = 1. For small *r*, a value of *g*(*r*) below 1 indicates repulsion. Note that for large *r, g*(*r*) always takes the value 1. Compared to the PCF, which is normalized between [0, 1], the DRP is an unscaled density of points at a distance *r* from a typical point. Ignoring edge effects, the PCF at a specific distance is equivalent to the average DRP over all cells scaled by the point process intensity.

#### L-function

The L-function^46^ is a scaled version of Ripley’s K-function *K*(*r*). In the planar case the L-function is defined as 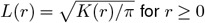 for *r ≥* 0. The function has a simple form (*L*(*r*) = *r*) in the case of the homogeneous Poisson process.

#### Comparison of summary functions for statistical testing

We note that the DRP is sensitive to binning parameters and edge effects. From a statical testing point of view, the L-function is superior to the K-function, the PCF and the DRP as it stabilizes variance and is taken as the norm for hypothesis testing in the spatial point process literature.^28^

### Pre-hoc cell type selection

#### Label noise filter

To reduce the label noise, we require the cell type to have sufficient number and intensity of high confidence cells, defined as cells with classification confidence score at least 0.7 (lower quartile of cell confidence scores) according to the reference clustering.^24^ For Glutamatergic cell types, we require the type to take at least 1% – average abundance in the presence of ∼100 cell types – of the total number of Glutamatergic cells of high confidence in the section of interest. For GABAergic cells, we further require the number of cells to be greater than 10 because there are fewer GABAergic cells per section. For each cluster, we study the sections where it has sufficient number of high confidence cells. Different sections are treated as replicated samples.

#### Minimal intensity filter

To remove cell types with low intensities from further analysis, we compare the intensity of high-confidence cells within individual types, as assigned by the reference clustering,^24^ to that of hardcore processes with the same interaction range. For each candidate cell type, we take the 20% quantile of its nearest-neighbor distance distribution as its approximated interaction range *R*. This corresponds to allowing at most 20% noise in the distance distribution of the corresponding hardcore process. We use the hardcore process with low intensity as our comparison benchmark. (See “Benchmark process for the minimal intensity filter” below.) This corresponds to the hardcore process with an interaction range *R* = 0.1 and self potential *λ* = 25 (i.e., approximated intensity 15) on a unit square, which rejects CSR at the limit significance of *p* = 0.05 in the absence of independent, additive noise points. To make the benchmark intensity comparable to the cell type intensity at the individual interaction distances, we rescale the benchmark intensity by a factor of 1*/*(10*R*)^2^. Equivalently, we can rescale the candidate cell type intensity (rescaled intensity) by a factor of (10*R*)^2^.

We focus on regions of high density for each cell type and compare its intensity to the benchmark. To avoid edge effects and region bias, we require reasonable homogeneity and sufficient area of the region under consideration. Specifically, we require that the area of the region should be at least (10*R*)^2^, comparable to the benchmark window size adjusted for *R*. We implement this using kernel smoothing as follows. We first calculate the maximum density across the section of interest. Then, we grow the window size by decreasing a density threshold (starting from the maximum density) until the area of the region meets the area constraint. We reject the section for that cell type if the resulting density no longer satisfies the intensity threshold. Hence, we use this intensity filter independently for each section of each candidate cell type, and exclude those whose intensity falls below the threshold.

#### Secondary label noise filter

For each cluster that passes through the intensity filter, we calculate the ratio of the number of high confidence cells to the total number of cells of the cluster (HC ratio) in every qualified section.

Considering that low HC ratios are indicative of label noise, we further exclude those cell types whose qualified sections all have low HC ratios (≤ 0.5), i.e., cells with confidence score greater or equal to 0.7 only take less than 50% of the total cells in the section. Importantly, the threshold is chosen such that HC= 0.5 corresponds to the noise/label ratio (Theorem 1) *α* = 1, where even the hardcore process with high intensity decays to a CSR. We note that more restrictive parameters can be chosen and our filters are robust to different parameter choices.

#### Summary of filtering steps

Supplementary Table S2 shows the filtering results for Glutamatergic and GABAergic clusters. There are around 100 Glutamatergic and GABAergic clusters in each section, 10-20 of which have a sufficient number of high confidence cells. The intensity filter further excludes around 50% of the candidate cell types in each section. In total, there are 18 Glutamatergic cell types and 10 GABAergic cell types that are qualified in at least one section. We show the HC ratio for each qualified cell type in each section (Supplementary figure S2).

### Benchmark process for the minimal intensity filter

We use benchmark simulations to identify a minimal intensity below which the power of statistical testing is not enough to detect structure in the point process. For this, we take the unit square as our study region and fix the hardcore interaction range *R* = 0.1. We note that this does not affect the generality of our study as rescaling is always possible: if we consider a different interaction range, we can choose a different window size and rescale the point pattern to window of unit size.

We simulate Strauss hardcore point patterns of high, medium and low intensities, such that the self-potential *λ* is equal to 100, 50 and 25, respectively. The corresponding true intensity is equal to 34, 25 and 15 approximately, using Poisson-saddlepoint approximation.^47^ The generated instances have 32, 25 and 15 points (Supplementary Figure S3). We note that, if we approximate the coverage region of each point by a circle of radius *R* = 0.1, then the total coverage of the points is approximately 1 in the high intensity case, the area of the unit square, ignoring edge effects.

The noise points *Y* are assumed to follow a Poisson process throughout the study region. Since we are interested in the noise/label ratio, we condition on the number of noise points in the study region. We define the label noise ratio *α* to be the number of the noise points divided by the number of points in the baseline patterns. We gradually add noise points to the baseline point patterns, from zero to the equal number of points as the baseline. We would anticipate the mixture process to show a gradual decay to the completely spatial randomness (CSR).

We quantify the approach to CSR using the DCLF hypothesis test based on the *L*-function. We take 99 CSR point patterns at each noise proportion *α* (See Theorem 1) for a starting and incremental p-value of 0.01. To stabilize the outcome, we fix the reference CSR patterns to be the same for each *α* ratio.

We plot the results for the three cases in Supplementary Figure S4. First, in all cases, the increase in p-values is significant. This confirms our intuition that the mixture process gradually decays to CSR. Moreover, we note the significance level of 0.05 corresponds to different levels of noise proportion *α* for different cases. Specifically, it corresponds to *α* = 0.4 in high intensity case, *α* = 0.2 in medium intensity case, and around 0 in low intensity case. The implies that the pattern is more robust to noise as we increase intensities of the baseline hardcore process. Moreover, the ability to detect the low-intensity Strauss pattern disappears as soon as any noisy points are added.

We further explore the effect of the point process intensity by testing the p-value under the DCLF test for hardcore process realizations with different intensities. Specifically, we consider hardcore processes with intensity parameters ranging from 10 to 50. For each setting, we simulate 10 point processes and test for statistical significance for each of them. We take the average over these 10 samples and plot the corresponding results in Supplementary Figure S5, which shows that the p-value decreases with the intensity parameter. It starts at around *p* = 0.2 when the intensity parameter (self-potential) is 10, implying insignificance, and decreases to *p* = 0.05 at around 25, the intensity parameter corresponding to the low intensity case studied above (Supplementary Figure S4). It then stabilizes below 0.05 for intensity parameters above 40. We thus take the low intensity case with a self-potential of 25 as our benchmark intensity for cell type selection.

### Selection of regions of interest

We perform statistical testing on tightly selected regions of interest to avoid assigning false significance to the fact that cells of a given type typically do not cover the whole brain. To avoid bias in region selection, we consider all cells of a given cell type, not just the cells annotated as “high-confidence”. For clusters and supertypes, we begin by considering anatomical parcellation substructures with non-zero counts in any section passing the intensity filter for that type as the initial window. For GABAergic cell types, this often corresponds to the whole section because GABAergic cells of a given type typically spread over the whole cortical section. For cell types with layer specification (e.g., L2/3 IT CTX Glut_1), we use all cells in the subclass they belong to (e.g., L2/3 IT) to identify the relevant parcellation substructures. We define the initial observation window for each cell type and section as the union of parcellation substructures with above-average abundance and above-median intensity per section. Hence, we include every parcellation substructure with reasonable density for the cell type of interest in each section.

We refine the initial observation window to relatively homogeneous regions using a kernel smoothing method. Specifically, we convolve the point pattern with an isotropic Gaussian kernel to smoothen the point pattern. We choose the standard deviation (bandwidth) of the kernel as half of the value suggested by likelihood cross-validation suggested in Ref.^48^ Following kernel smoothing, we select regions with above-average intensity and erode the selected region using the bandwidth of the kernel as the erosion parameter to remove the spurious regions introduced by smoothing.

Projecting a slice of 3D tissue onto a 2D surface ignores the thickness of the section (10*µm*) and distorts the distances. This distortion can produce spurious spatial clustering for some cell types, especially the Glutamatergic clusters which are more abundant than the GABAergic ones in the cortex. To alleviate this issue, for Glutamatergic clusters with nearest neighbor distances less than 10*µm* (Table 1), we further impose an upper limit on the intensity of the selected region at two standard deviations above the average intensities.

Supplementary Table S3 shows the corresponding statistics for Glutamatergic clusters before this extra step of region selection. Importantly, we note that the two clusters that fail to reject CSR, 0453 L6 CT CTX Glut 5 and 0004 CLA-EPd-CTX Car3 Glut 1, have higher values for both average percentiles (of nearest neighbor distances less than the section thickness) and point process intensities before this extra step of region selection (Table S3).

### Simulation of 3D soma locations under the spatial randomness hypothesis

The dataset used in this study^25^ comprises 2D images of 10*µm*-thick sections with 200*µm* gaps between consecutive sections. The cells in the same section are thus assumed to be on the same surface, distorting the actual distances between the cells. For instance, cells may appear to be right on top of each other in the 2D section, destroying the underlying hardcore process. We note that this is more of a concern for Glutamatergic clusters as the Glutamatergic cells are more numerous compared to GABAergic cells in the cortex.

As a first step towards simulating soma locations under the spatial randomness hypothesis, we obtain an unbiased estimate of the distribution of soma sizes from a volume election microscopy dataset.^30^ We examine the effect of 3D to 2D projection by studying section cuts from Strauss hardcore processes with hardcore distances specified by estimates of the soma radius.

To simulate a section, we consider a cuboid of size 200*µm* × 200*µm* × 50*µm* (width × length × height). We define the benchmark as cells being separated by hardcore distances only. We consider hardcore interaction ranges corresponding to both the minimum (7.4*µm*) and average soma diameters (10*µm*) based on the electron microscopy dataset and simulate 10 point patterns that respect the hardcore constraint. We observe that projecting the section to 2D completely destroys the hardcore distance of 7.4*µm* while the hardcore distance of 10*µm* is decreased to *∼* 5*µm* (Supplementary figure S12).

We further explore the effect of projection by studying 3D point patterns of different intensities. Intuitively, as the intensity of the hardcore point pattern decreases, the hardcore distances will become less significant and the process will approach complete spatial randomness. We simulate 3D Strauss hardcore processes of different intensities with the measured minimum soma diameter as the hardcore distance. For each 3D point pattern, we calculate the intensity and p-value of the projected 2D pattern using the 1-sided DCLF test at the specified interaction range. We take 99 CSR point patterns for a starting and incremental p-value of 0.01. Supplementary Figure S12 shows that the projected point patterns become statistically indistinguishable from CSR as intensity drops below 1500 cells/mm^2^.

In summary, results for clusters with large 3D projection errors should be carefully interpreted to avoid falsely declaring a clustering effect due to the apparent overlap of cells. Moreover, statistically significant results obtained by our 2D testing framework for clusters with small interaction ranges and low intensities compared to the benchmark process above cannot be solely attributed to hardcore separation (due to the physical space the soma occupies). Instead, other factors, such as softcore repulsion with a longer interaction range, should be present.

### Testing of spatial randomness for cell populations

We consider the DCLF-test based on the *L*-function by comparing the theoretical *L*-function to the *L*-function of the observed point patterns of interests. For GABAergic cell types, we use the Strauss hardcore process with a hardcore distance of 10*µm* (average soma diameter) as the theoretical benchmark of CSR because GABAergic clusters are less affected by the projection error. On the other hand, Glutamatergic cells are more abundant compared to GABAergic cells and suffer more from projection to 2D. As studied in the previous section, the hardcore distances could be destroyed due to projection so that the Strauss hardcore process is not suitable to serve as the CSR benchmark. Therefore, we use the homogeneous Poisson process as the CSR benchmark for Glutamatergic cell types.

For each sample population, we conduct the 1-sided DCLF test based on *L*-functions of 199 CSR point patterns with the same number of points as the observed point pattern in the observation window. The DCLF test is sensitive to the interaction range chosen for the test. We consider distances from the hardcore soma separation (10*µm*) to an upper limit such that interactions beyond that limit are not of interest anymore. We set the upper limit as the median nearest neighborhood distance across sections where the cluster passes the noise and minimum intensity filters.

We calculate the minimum p-values at which we reject CSR for regularity over a range of interaction distances (every 1 *µ*m from the hardcore separation distance to the upper limit). The p-values corresponding to individual sections are combined using Fisher’s method and corrected for multiple testing using the Benjamini-Hochberg procedure. To stabilize the outcome, we fix the reference CSR patterns to be the same for all distances. If the corrected p-value is below 0.05, we say the point pattern *rejects CSR*.

### Effective cluster count

Consider a population 𝒳 in a study region *W* that consists of *N* unique cluster labels, i.e, *{c*_1_, …, *c*_*N*_ *}*. For each cluster *c*_*i*_, we calculate its relative abundance *p*_*i*_ = *N* (*c*_*i*_|𝒳)*/*Σ_*j*_ *N* (*c*_*j*_|𝒳) where *N* (*c*_*i*_|𝒳) is the number of samples of cluster *c*_*i*_ inside the study region *W*. We define the effective cluster count (ECC) of 𝒳 in *W* as

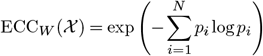

If the cluster is studied on multiple brain sections *k* = 1,…, *K*, we calculate its ECC for study region *W*_*k*_ in each section *k* and take a simple average over all sections, i.e,

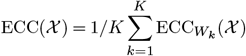

The definition of ECC relates to the concept of types ((49), Section 11.1) in information theory. We can treat the population 𝒳 in study region *W* as a marked point process of *n* points where the mark of each point corresponds to its cluster label *c* ∈ {*c*_1_, …, *c*_*N*_}. We then tabulate the point process in a study window *W* as a sequence of *n* symbols from an alphabet *A* = {*c*_1_, …, *c*_*N*_} (arbitrary ordering). We denote the tabulated sequence as ***x*** = (*x*_1_, … *x*_*n*_). The type *P*_***x***_ of the sequence ***x*** is defined as the relative proportion of occurrences of each symbol of *A*, which corresponds to the relative abundance of each cluster.

Note that the type does not uniquely define a sequence. Let *P*_*n*_ denote the set of sequences of length *n*. If *P ∈ P*_*n*_, the type class of *P* is defined as *T* (*P*) = *{****x*** *∈ A*^*n*^|*P*_***x***_ = *P}*. By Theorem 11.1.3 in (49), for any *P ∈ P*_*n*_,

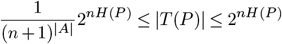

where the entropy is measured in bits. The theorem states that there are an exponential number of sequences of each type, i.e, |*T* (*P*)| =. 2^*nH*(*P*)^.

Now we introduce the concept of effective count by taking the *n*-th root of the size upper limit and using nats as our entropy measure. Note that the size of *T* (*P*) is the number of ways of arranging *np*_1_, … *np*_*N*_ objects in the sequence and there are *n* elements in total. If we assume there are effectively only *Ñ* symbols in the sequence, each component can only take *Ñ* values, so there are at most *Ñ*^*n*^ choices for arrangement. By equating the number of arrangements, we have

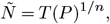

defined as the ECC of population *X* in study region *W*.

Moreover, we assume that different brain sections are replicated samples of the cell type distribution. Therefore, the types in different study regions can be thought of as i.i.d. samples form the type distribution. We can then take a simple average over multiple brain sections to obtain the overall ECC of the cell type *𝒞*.

## Supplementary Notes

### A. Distribution of the radii of cell bodies

We obtain a distribution of soma sizes by studying soma segmentations from multiple areas of mouse visual cortex in the MICrONS dataset.^30^ The dataset consists of 94014 cells with extracted somatic measurements, including nucleus area and volume, nuclear folding metrics, soma area and volume and somatic synapse information. Each cell also has a nucleus id, soma id and a predicted cell type.

We approximate the soma by a sphere and estimate the soma radius from the soma volume measurement. Figure S11 shows distributions of the estimated soma radius for Glutamatergic and GABAergic cells.

Except for 5P-ET cells, the approximated soma radius is similar across Glutamatergic types. The minimum radius is ∼ 3.7*µm*, and the average is 5*µm*. The soma radius estimate for 5P-ET cells is significantly larger with a mean of ∼ 7*µm*. The soma radius estimate for GABAergic cells varies by subpopulation. The minimum radius estimate is 3.5*µm*, slightly smaller than the minimum estimate for Glutamatergic cells.

### B. Pre-hoc filtering steps

Figure S2 illustrates our filtering process. We demonstrate the minimal intensity filter (using 0645 Vip), the secondary label noise filter (using 0436 L6 CT), and the observation window selection process (using 0741 Pvalb). Table S1 lists the number of cell types selected by each filter in individual sections.

#### Label noise filter

In section 22, the cell type 0436 L6 CT has a high proportion of cells with confidence below 0.7 (black dots). As shown in Figure S2a, low-confidence (black) cells occupy a large proportion of the region of interest. The same happens for other sections of this cell type. Therefore, this cell type is filtered out from downstream analysis.

More broadly, cell type assignments and the confidence in these assignments in new spatial transcriptomic datasets (e.g., based on Refs.^24, 25^) are prone to batch effects, which should be removed prior to applying the label transfer code in Ref.^25^ or other similar software such as Allen Institute’s *MapMyCells* tool at https://knowledge.brain-map.org/mapmycells/process/.

#### Intensity filter

The “0645 Vip” cell type fails to pass through the intensity filter in section 22 (Table S2). The intensity of the high-confidence cells is low (Figure S2b) and visually comparable to the low intensity hardcore process simulation in Figure S3.

#### Observation window selection

As discussed in the Methods section, we perform statistical testing on tightly selected regions of interest. We use the “0741 Pvalb” cell type in section 24 for illustration. Firstly, we consider anatomical parcellation substructures where the cell type abundantly exists with non-zero counts. As shown in Figure S2c, we use all cells in the selected anatomical parcellation substructures to draw a boundary for the cell type 0741 Pvalb. This window is then intersected with the convex hull of 0741 Pvalb cells, giving the initial window. Then we refine the initial observation window using kernel smoothing method, giving the region of interest for 0741 Pvalb in section 24.

### C. Mixture point process theory

#### Mixture of Strauss and Poisson point processes

We seek to characterize a mixture of a point pattern corresponding to a pure cell type, modeled by the Strauss process, and a point pattern corresponding to cells that are incorrectly assigned to that same type (even though they actually belong to multiple different cell types), modeled by the Poisson process. The theorem below formalizes this setup and Figures S4, S5 illustrate the theorem for the special case of a hardcore process.

#### Edge effects

Point process statistics change at the boundaries of finite regions of interest. For instance, trivially, the interaction strength is not well defined beyond the limits of the observation window. In the formal analysis below, we will “ignore edge effects”. In practice, the edge effects can be corrected using various methods such as border correction, which focuses on an interior subregion of the observation window farther from the boundary than the interaction range. Moreover, the edge effects are negligible in the limiting cases of the self-potential *β* → ∞ in a finite observation window or the observation window size |*W*| → ∞ (in all dimensions) with a finite self-potential.

##### Theorem 1

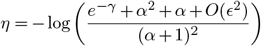

for some small *ϵ >* 0 with *ϵ* = (1 *−e*^*−γ*^)*/g*_1_, *g*_1_ *≥* 0.

*Proof:* Considering the mixture process as a pairwise interaction between two independent marked point processes, i.e. points belong to either the Poisson ***x***_**0**_ or Strauss ***x***_**1**_ processes, the mixture defines a *bivariate Strauss process*.^50^ The conditional intensity of this mixture process is

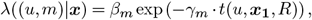

where *m* ∈ {0, 1}, *β*_0_ = *aβ, β*_1_ = *β, γ*_0_ = 0, *γ*_1_ = *γ*. When the mixture process is considered as an unmarked point process, the conditional intensity can be written as

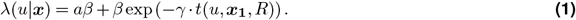

The conditional intensity of a Strauss process with an interaction strength *η*, self-potential *β* and interaction range *R* for the same mixture would be

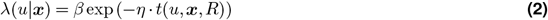

Solving Equations 1 and 2 for *η* reveals that the conditional intensity of the mixture can be put into the form of a Strauss conditional intensity:

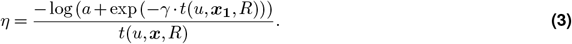

Since *a >* 0 and *γ ≥* 0,

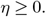

Moreover, since *α >* 0 and *t*(*u*, ***x***_**1**_, *R*) *≤ t*(*u*, ***x***, *R*),

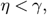

implying a weaker interaction strength.

While the expression for the interaction strength *η* in Equation 3 depends on the process ***x***_**1**_ as well as ***x***, there is no underlying inhomogeneity in either of the non-interacting, independent mixture components. Therefore, ignoring edge effects, *η* = *η*(*R*) and the mixture defines a valid Strauss process. To obtain such an expression, let *n*_0_ and *n*_1_ be fixed and *α* = *n*_0_*/n*_1_. The *K*-function is defined as

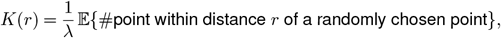

where *λ* is the intensity of ***x***_**0**_ ∪ ***x***_**1**_ (Section 4.3, (28)). Ignoring edge effects, the empirical *K* function is given by (Section 13.4, (51)),

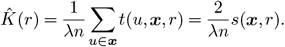

Assuming the interaction strength *γ* is fairly weak,^50^ we write

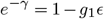

for some small *ϵ >* 0 and *g*_1_ is a non-negative constant. Since *η < γ*, we can write *e*^*−η*^ = 1 − *g*_2_*ϵ* where *g*_2_ *< g*_1_ is also a non-negative constant.

The K-function for the Strauss process with interaction strength *η* can be approximated to first-order in the interaction strength^50^ as

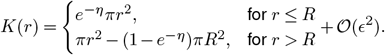

In this way, we can approximate *s*(***x***, *r*), the number of unordered pairs within a distance *r ≤ R* in ***x*** as

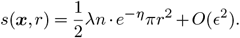

Similarly, for the Strauss component ***x***_**1**_, the number of unordered pairs within a distance *r ≤ R* is

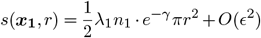

where *λ*_1_ is the intensity of ***x***_**1**_.

The *K*-function has a simple form for the Poisson point process: *K*(*r*) = *πr*^2^, and the number of *r*-close pairs is given by

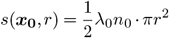

where *λ*_0_ is the intensity of ***x***_**0**_.

To compute the number of *r*-close pair in the mixture, we consider the cross-type *K*-function between the two components ***x***_**0**_ and ***x***_**1**_. Since ***x***_**0**_ and ***x***_**1**_ are independent processes and the mixture process results from a set-theoretic union of ***x***_**0**_ and ***x***_**1**_, the cross-type *K*-function for the random superposition model (Section 5.3, (28)) is given by

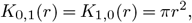

Where

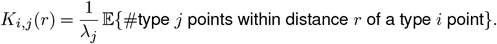

If we denote the *r*-close pairs of component ***x***_**0**_ and ***x***_**1**_ as *s*(***x***_**0**_, ***x***_**1**_, *r*), i.e.,

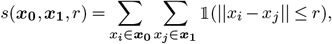

we have

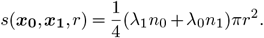

Since ***x***_**0**_ and ***x***_**1**_ are point patterns inside the same region *W, λ* = *n/*|*W*|, *λ*_0_ = *n*_0_*/*|*W*| and *λ*_1_ = *n*_1_*/*|*W*|. Noting that

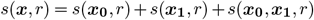

for *r ≤ R*, we obtain an approximate explicit expression for the interaction parameter *η* as

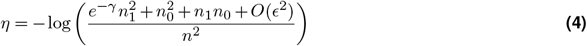

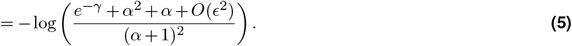

##### Remark

: Clearly, when *α* = 0, *η* = *γ* and as *α* → ∞, *η* → 0. When the baseline process has large interaction strength s.t. *γ* ≫ 1 and *α* = 1, *η* is approximately log(2).

### D. Statistical testing for individual cluster-section pairs and its aggregation

In the main text, we report raw p-values aggregated over sections according to the Fisher’s method. In this section, we show the p-values for individual sections and adjusted p-values using the Benjamini-Hochberg procedure for all clusters of interest (Tables S4, S5, S7, S8).

For each type and the section where it passes the noise and minimum intensity filters, we conduct the 1-sided DCLF test with 5 replicates using different seeds. The DCLF test covers the interaction range from the hardcore separation of 10*µm* to the range defined by the median value of nearest neighborhood distributions. The minimum p-values over the interaction range are recorded per replicate and averaged. Then the aggregated p-value is calculated according to Fisher’s method and corrected using the Benjamini-Hochberg procedure.

### E. Example case studies

#### Simulated patterns for exemplar cell types

In Figure S1, we show two more simulated patterns for the exemplar cell types shown in Figure 1.

#### Pvalb pair 0740 and 0742

We study the mixture of Pvalb 0740 and Pvalb 0742 as a single cluster. Table S9 lists the number of cells of each of these two types in different cortical regions. Since the corresponding set of points does not cover the entire section (Figure S17a), we define the initial observation window for this mixture pair as the union of the observation windows for the individual transcriptomic types.

Figure S17b1 shows the nearest-neighbor distribution of the two types as a single cluster and demonstrates that the cells can get arbitrarily close to each other (*p*-value for CSR: 0.99, 1-sided DCLF test), spatially cluster (weakly) in distances smaller than 50 microns, then their curve closely follows that of CSR. Moreover, we examine the pairwise correlation function for the mixture pair in Figure S17b2. Similarly, the pair correlation function plot is similar to that of the Pvalb subclass (no hardcore distance and no significant local repulsion). Intuitively, for the point process of each cluster in the observation window, the other one serves as noise that destroys its homotypic avoidance property. We further study their interaction by treating them as a marked point process inside the observation window where the marks are cluster types. Unsurprisingly, Figure S17b3 and Figure S17b4 shows that the two types appear to be spatially clustered at close distances (∼50*µm*) and repel each other at long distances, at the scale of cortical parcellations (Table S9).

#### Vip supertype 0173 and cluster 0624

As noted in the main text, the supertype 0173 Vip is an exception of the tree hierarchy diagram. It has an ECC value of 2.54, although it displays significant repulsion (one-sided DCLF test, *p* = 0.013). We examine this supertype and its child cluster 0624 Vip in section 24.

Figure S18a shows the spatial distribution of supertype 0173 Vip neurons in sagittal section 24. For the supertype and its dominant cluster 0624 Vip (blue dots), we plot the corresponding windows for analysis (supertype: black outline, cluster: blue outline). The selected region for the supertype 0173 includes the selected region for the cluster 0624. The region for supertype 0173 is mainly covered by that of 0624, with uncovered regions consisting of 2 cells from 0623 Vip and 4 cells from 0625 Vip.

Figure S18b shows the L-functions and pairwise correlation functions for the supertype 0173 and cluster 0624, which demonstrates similarities between the supertype and the cluster in terms of the hardcore distance and the range of local repulsion. Since both cell types 0623 and 0625 have few samples in this observation window, and do not significantly disturb the homotypic avoidance behavior of 0624 Vip, we hypothesize that an oversplit in the reference hierarchy may contribute to the significance at this Vip supertype level.

### F. Pairwise correlation functions for all cell types

Table S4 and S5 show that inhibitory clusters generally have higher p-values per section compared to excitatory clusters. That is, inhibitory clusters have less significant deviations from CSR in general, based on the proposed testing paradigm. This is also hinted by Figure 1D, where the red line (observed L-function for 0109 L2/3) deviates more from the shaded upper and lower envelopes of the CSR simulations, compared to the blue line (observed L-function for 0741 Pvalb).

To further explore this difference between excitatory and inhibitory clusters, we plot the pairwise correlation functions of the cell bodies in every cluster that reject CSR and has at least 10 samples in a window of interest (Supplementary Figures S13 and S14). For each cluster, we pool the PCF over selected sections with *p <* 0.05 and plot over the range from 0*µm* to 50*µm* for excitatory clusters and to 120*µm* for inhibitory clusters (maximum of `‘R’` in Table 1). We use the standard kernel estimator to compute the PCF by default.^36^ However, for clusters with non-zero average percentiles (`‘%’` in Table 1)), we use a modified estimator^51^ to improve the distortion near zero due to the 3D projection error (Figure S12). We note that PCF is highly unstable at near-zero distances compared to the L-function, with or without the modified estimator. For this reason, we choose L-functions for statistical tests over PCFs.

In *absolute* terms, inhibitory clusters generally show clear hardcore distances and have much longer (2-3 times) radius of homotypic avoidance compared to excitatory clusters. However, we note that PCFs for most of the inhibitory clusters cross the value of 1 before its upper interaction range `‘R’`, while the PCFs for most of the excitatory clusters stay below the value of 1 throughout the plotted range (excluding mild initial increases, presumably due to the projection error). Thus, excitatory clusters have longer *relative* interaction ranges and higher interaction strengths compared to inhibitory clusters (Tables 1, S7, and S3).

Figure S16 displays PCF plots for inhibitory subclasses, demonstrating clustering of somata for distances less than ∼ 20*µm*.^17, 18^

### G. Density recovery profiles of example populations

The PCF is closely related to the density recovery profile (DRP),^34^ which is often used in retina research.^34,3,21,35^ Compared to the PCF, which is an estimate of the probability of finding a pair of points a distance *r* apart, the DRP is an unscaled density of points at a distance *r* from a typical point. Ignoring edge effects, the PCF can be derived directly from the DRP by taking an average of DRP over every cell in the point process and normalizing by a factor of *λ*^2^ where *λ* is the point process intensity. Figure S15 shows the DRP for three selected types (two of them are given as examples in Figure 1), which demonstrates the similarity of the profiles obtained from cortical cell types to those obtained from retinal cell types. (See, for instance, Ref.,^34^ Figure 2A in Ref.,^3^ Figure 1c in Ref.,^21^ or Figure 1E in Ref.^35^) Figure S15 also shows cross-type DRPs for Pvalb 0740 and Pvalb 0742 in Section 23 (given as a case study in Figure S17b), and for Pvalb 0740 and Pvalb 0741 in Section 24. We observe that, compared to the DRP for 0741 Pvalb, the values increase at short distances, indicating that self-avoidance disappears when cell types merge. The cross-type DRPs also agree with Figure S16 and show that interneurons tend to cluster spatially at near distances at the subclass level.^17, 18^ We note that the DRP is calculated for each cell in the point process and can be subject to large variations, especially for cells at the boundary of the observation window. Moreover, the DRP is highly sensitive to binning width. Here, the bin widths are chosen manually so as to better compare with Ref.^3^

### H. Differential gene expressions for all cell types

Table S6 compiles the differentially expressed genes for each of the transcriptomic types studied in this work. All cell types pass pairwise differential expression gene analysis test that ensures that there are multiple strong markers that can be used to discriminate the cell types from each other. The table is derived from the full table at https://alleninstitute.github.io/abc_atlas_access/descriptions/WMB-taxonomy.html.

### I. Gallery of cell types and selected region of interest

Supplementary Figures S6, S7, S8, S9, and S10 show a gallery of soma (point) locations for each cell type with their selected region of interest. For GABAergic types, we plot the corresponding windows for analysis in blue outline. Recall that due to 3D to 2D projection issues, for the more populous Glutamatergic types, we adjusted the region selection step to impose an upper limit at two standard deviations above the cell type average, which was discussed in the Methods section. We show the regions before this extra step in red outline, and those due to the extra step in grey.

### J. Supplementary dataset (Coronal sections)

We further confirm our results in a second dataset by Ref.^24^ We take three coronal sections at the dorsal part of the mouse cortex (Section 59-61). Figure S19 illustrates the three brain sections we consider in the dataset. 25 Glutamatergic types and 5 GABAergic types pass through the intensity and label noise filter. We use the same hypothesis testing method for these cells types across brain sections. Table S10 shows the hypothesis testing result corrected for multiple testing. The results gives qualitatively similar conclusion as the sagittal dataset. Importantly, a majority of Glutamatergic clusters and all GABAergic types display homotypic avoidance.

## Supplementary Figures

**Figure S1.**
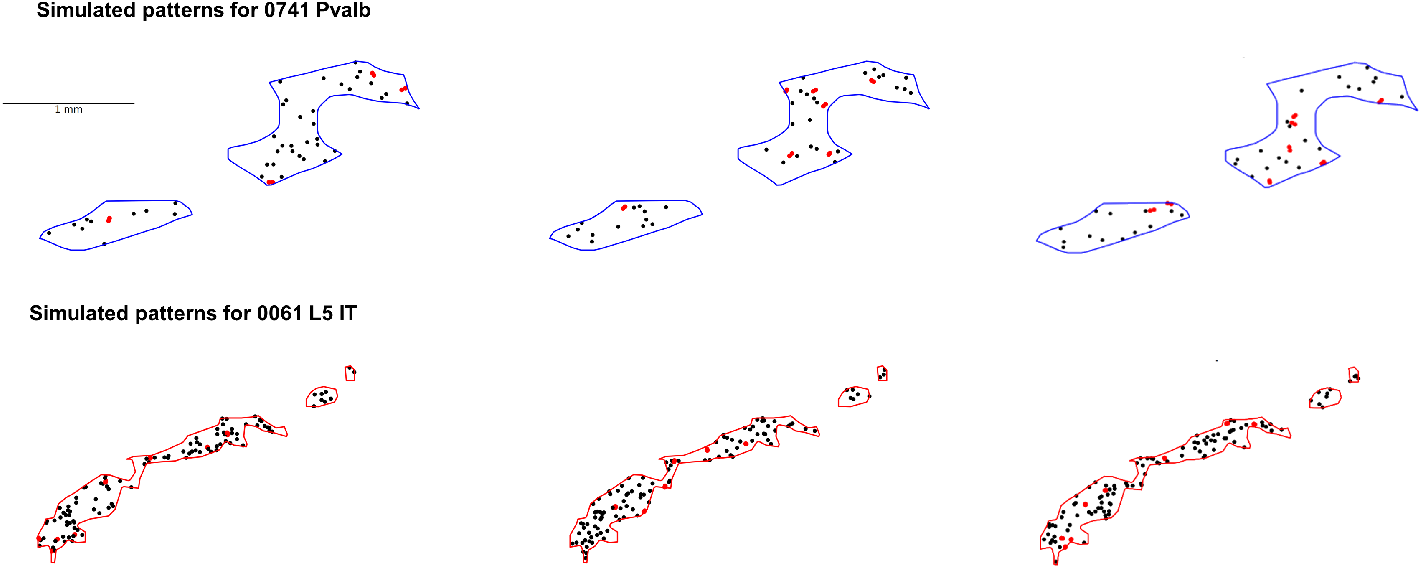
More simulated patterns for the cell types shown in Figure 1.

**Figure S2.**
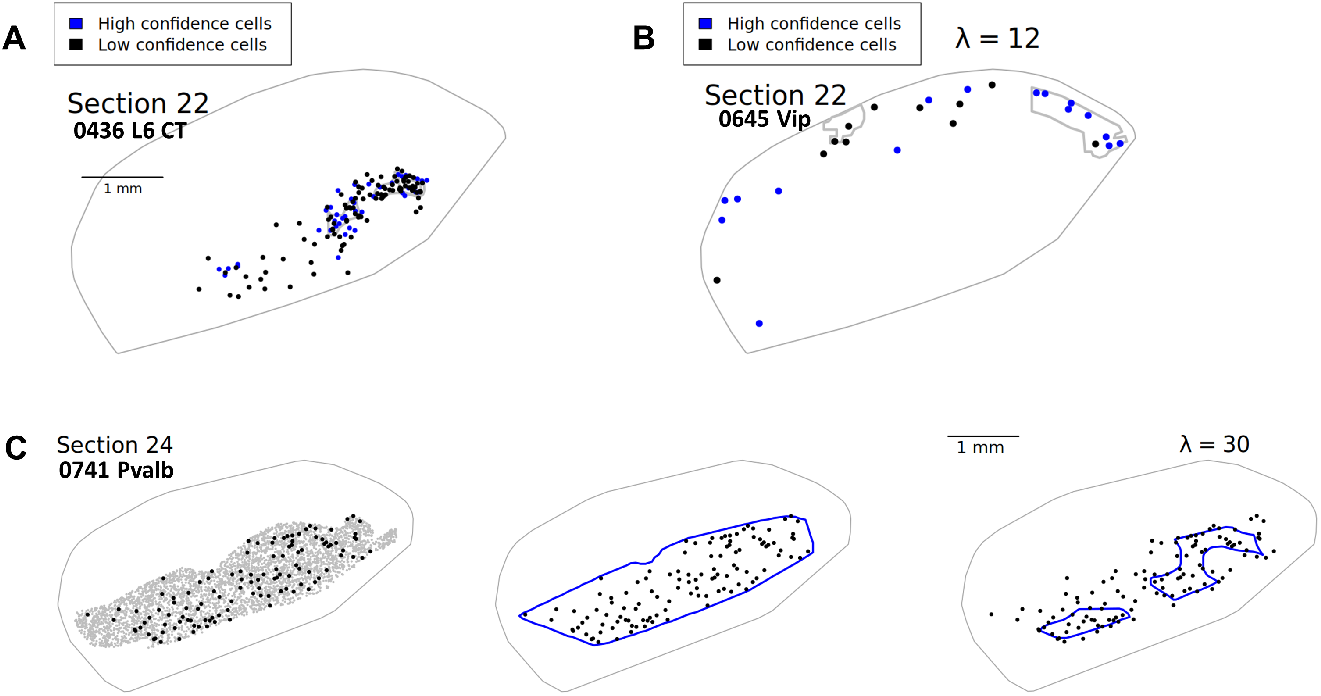
Graphical illustration of label noise filter, intensity filter and observation window selection. **A**. Label noise filter illustrated using 0436 L6 CT in section 22. **B**. Intensity filter illustrated using 0645 Vip in section 22. The intensity of the high-confidence cells is low (rescaled intensity *λ* = 12) and visually comparable to the low intensity hardcore process simulation (*λ* = 15) in Figure S3. **C**. Observation window selection illustrated using 0741 Pvalb in section 24 (rescaled intensity *λ* = 30, *p*-value for CSR: 0.007, 1-sided DCLF test). Black dots: 0741 Pvalb cells. Left: Use all cells in the selected anatomical parcellation substructures (grey) to draw a boundary. Middle: Initial window derived as the intersection of the grey boundary with the convex hull of 0741 Pvalb cells. Right: Refine the initial observation window using kernel smoothing method.

**Figure S3.**
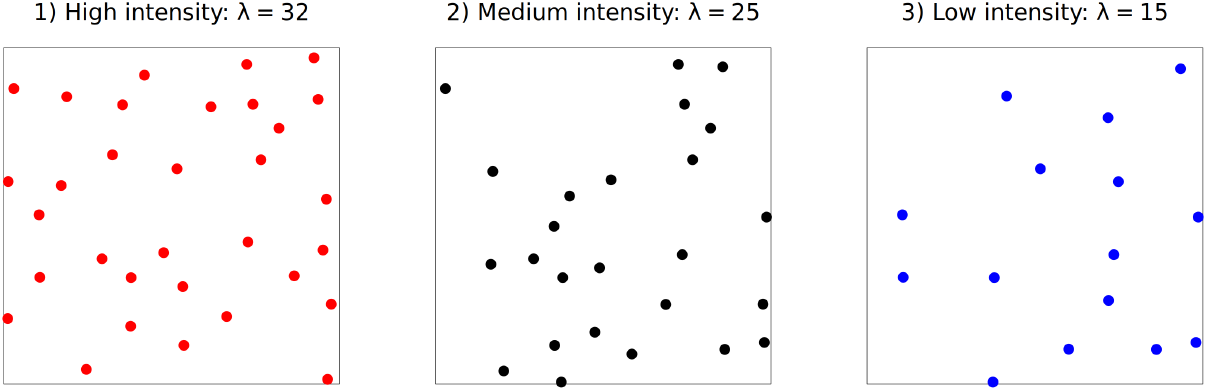
Sample Strauss hardcore process point patterns of high, medium, and low intensities.

**Figure S4.**
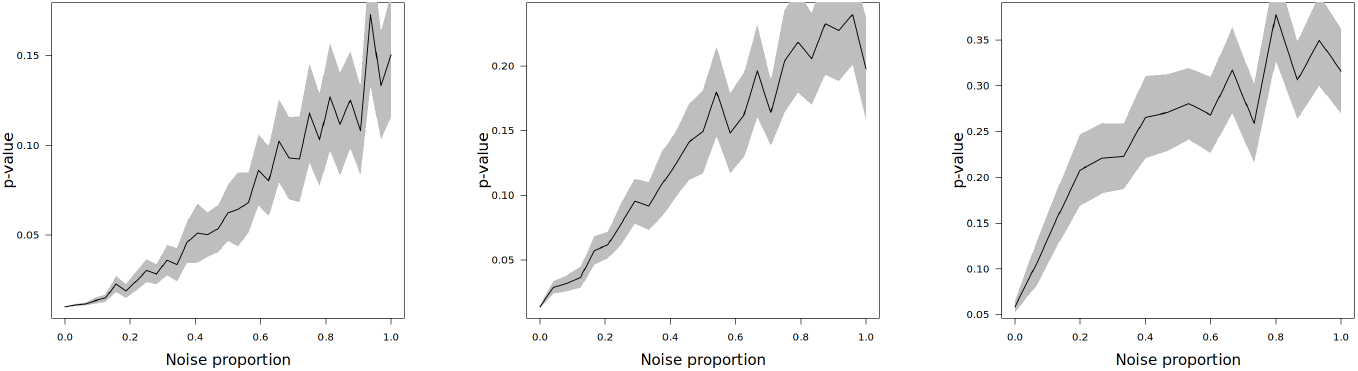
Deviation from CSR for mixtures starting from different hardcore processes. Left to Right: hardcore processes of high, medium and low intensities. In all cases, a point pattern generated from the Poisson point process is independently added to the point pattern of the hardcore process. Noise proportion denotes the ratio of Poisson points to hardcore points. *y*-axis reports the p-value obtained from the 1-sided DCLF test. We observe decrease in statistical significance with increase in noise proportion. The pattern becomes more robust to noise as the intensity of the baseline hardcore process increases.

**Figure S5.**
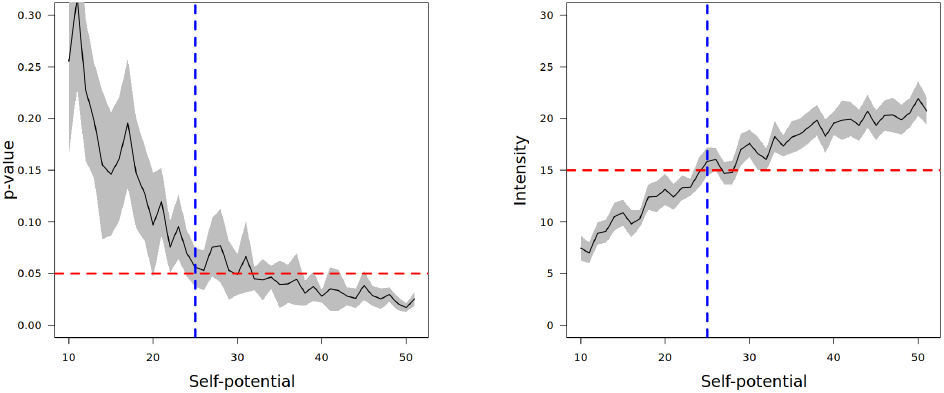
Effect of initial point process intensity. Left: Change in statistical significance with respect to intensity of the hardcore process. *y*-axis same as that of Figure S4. The p-value decreases to 0.05 at self-potential around 25, approximately corresponding to intensity around 15 (Right), which is taken as our benchmark intensity to be used for cell-type selection.

**Figure S6.**
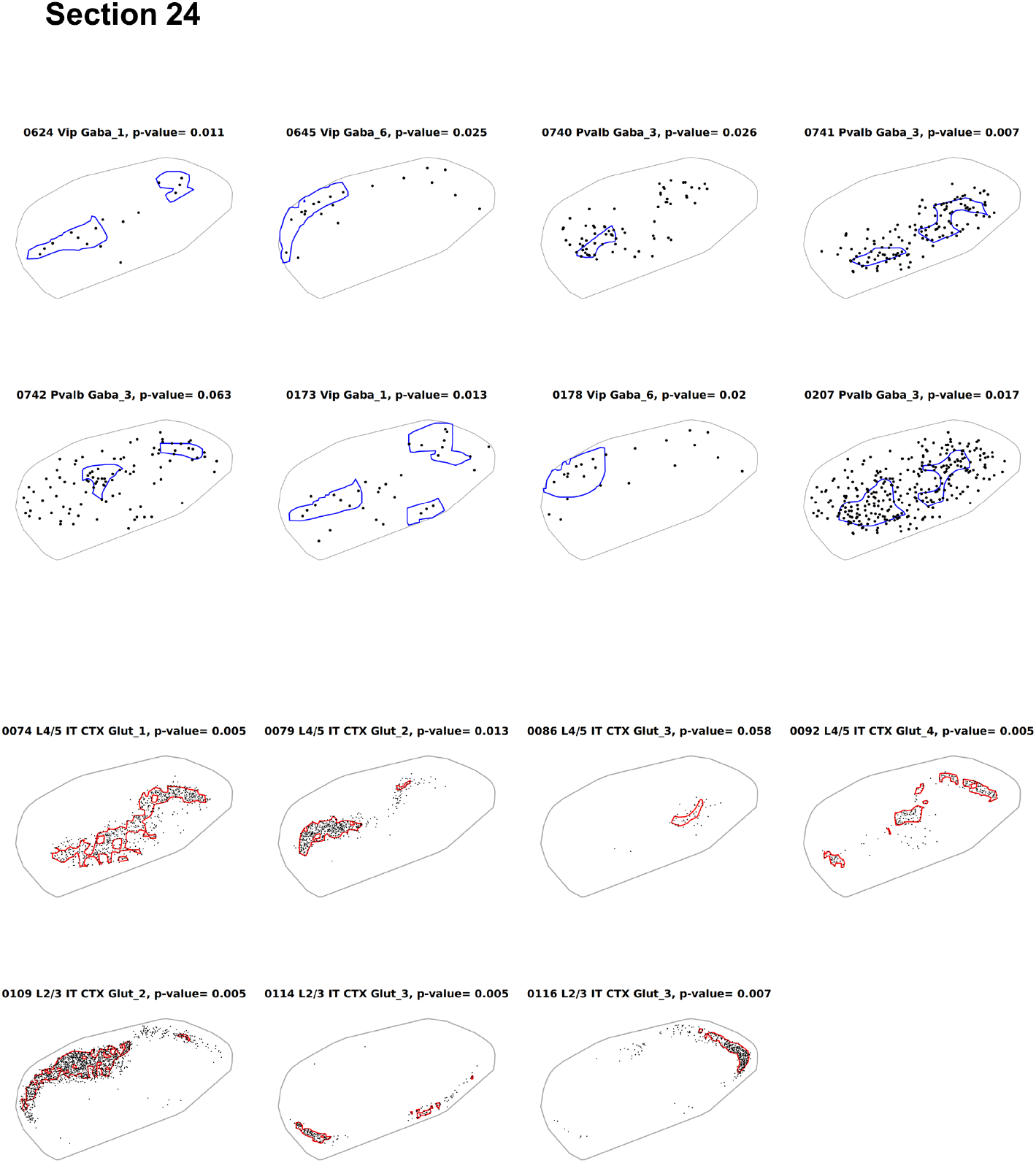
Soma locations and regions of interest for all studied cell types in Section 24.

**Figure S7.**
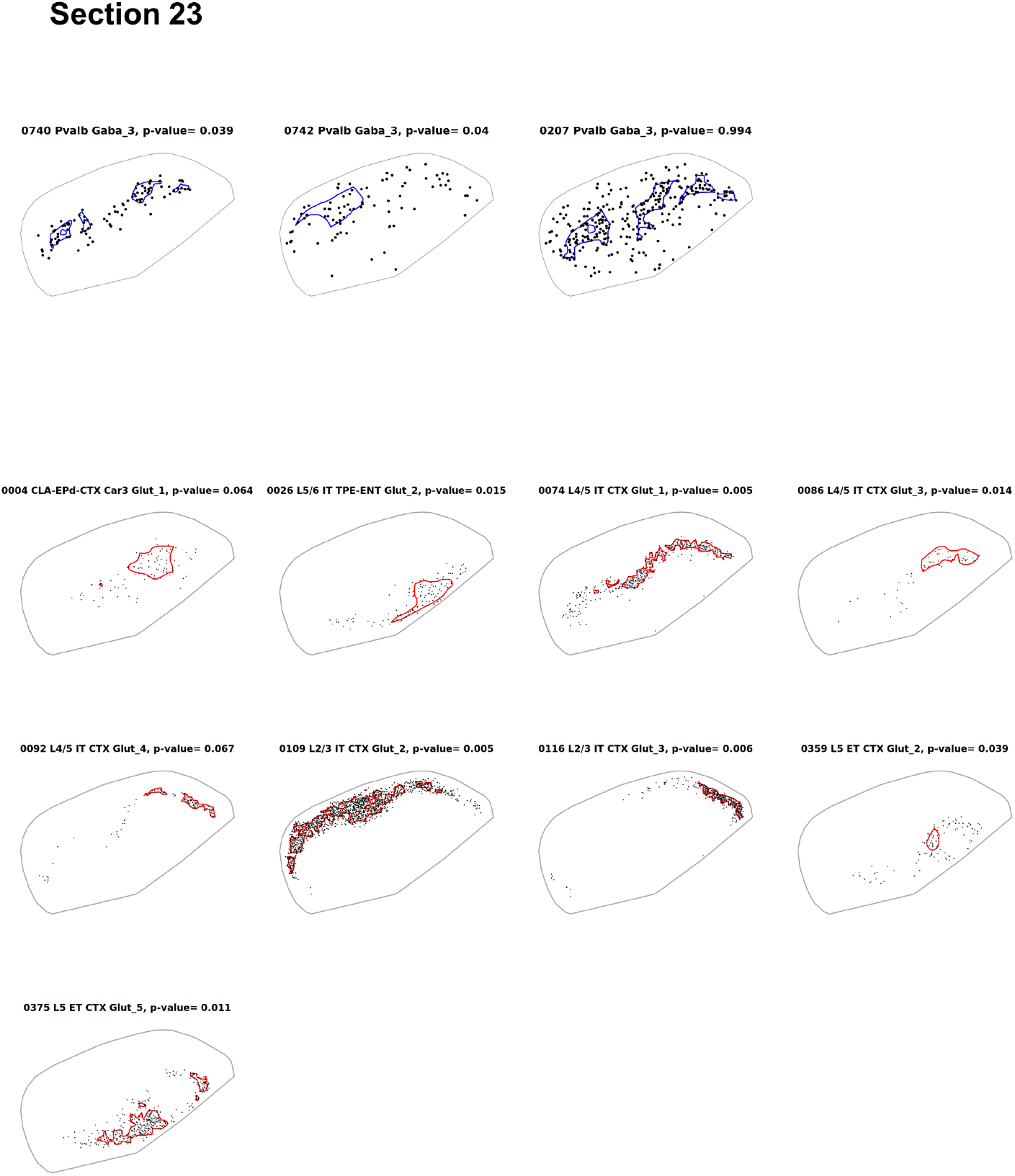
Soma locations and regions of interest for all studied cell types in Section 23.

**Figure S8.**
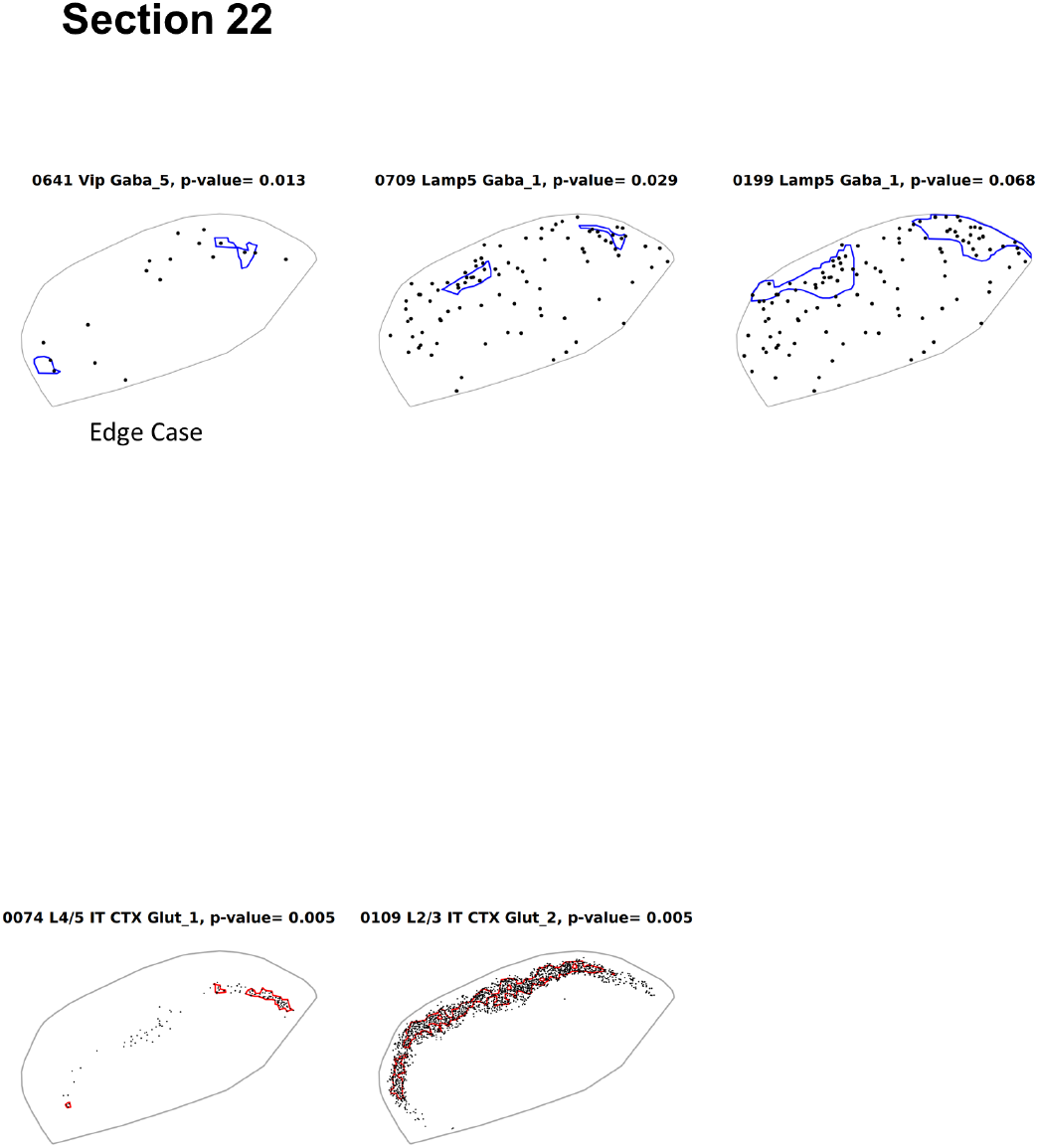
Soma locations and regions of interest for all studied cell types in Section 22.

**Figure S9.**
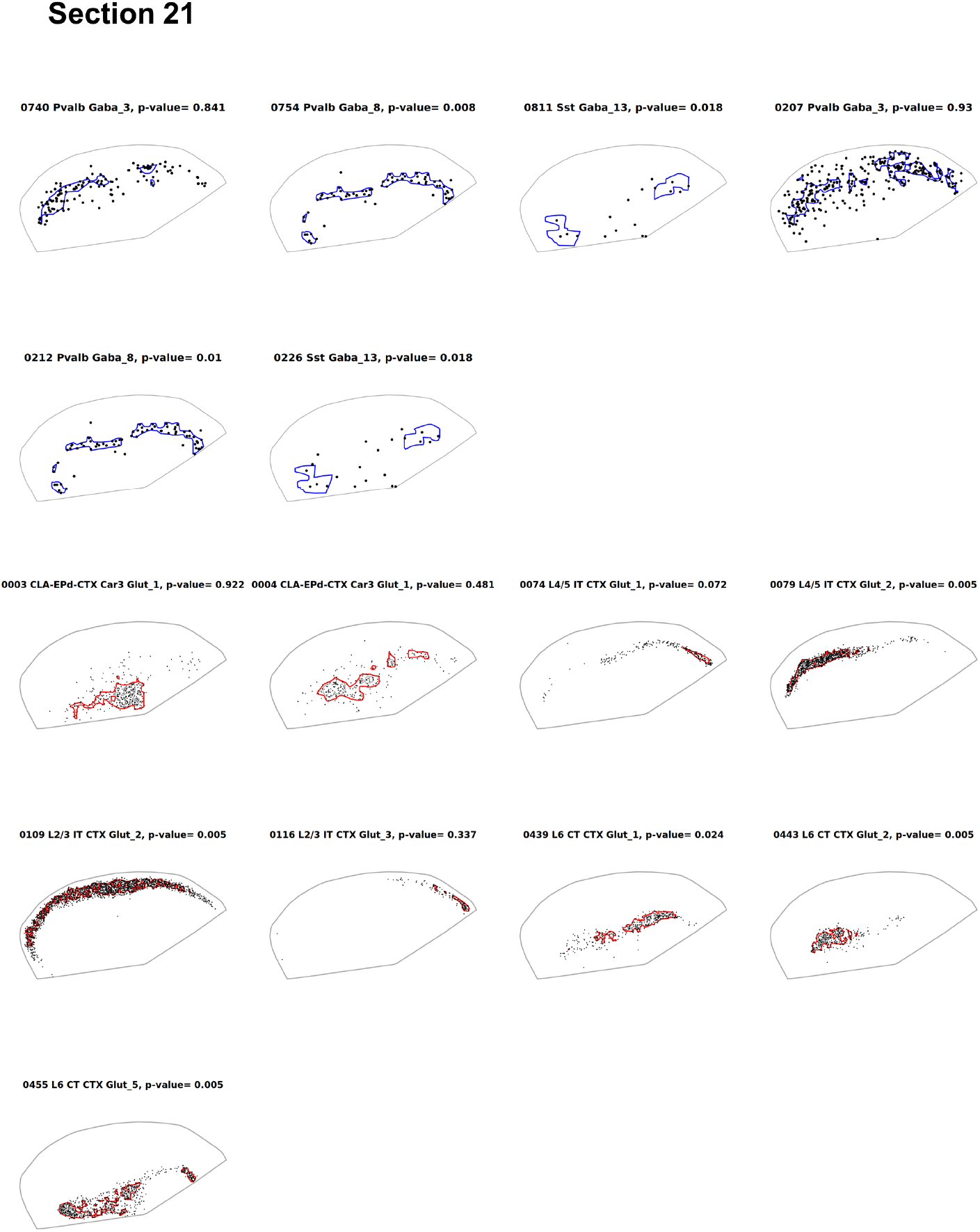
Soma locations and regions of interest for all studied cell types in Section 21.

**Figure S10.**
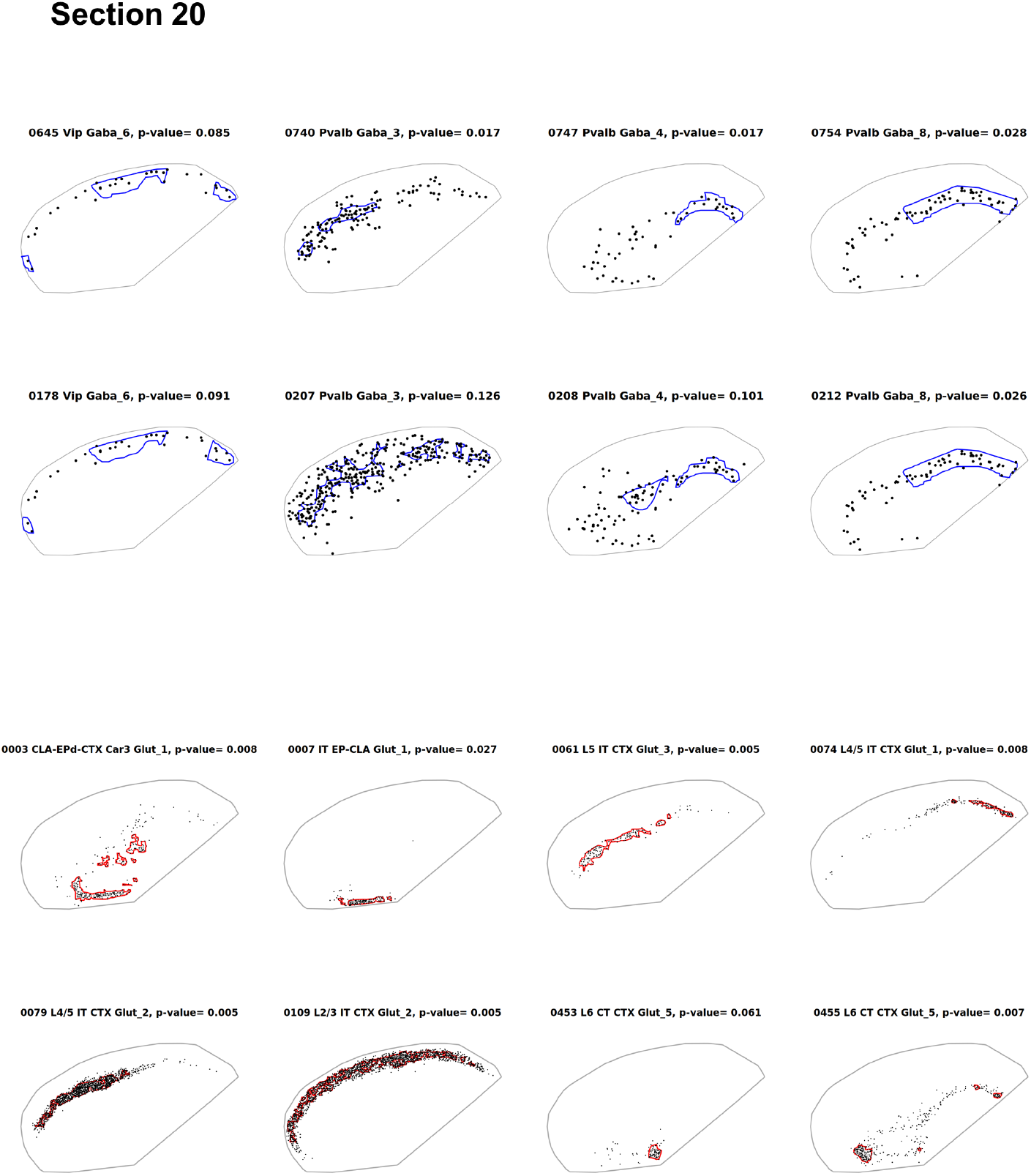
Soma locations and regions of interest for all studied cell types in Section 20.

**Figure S11.**
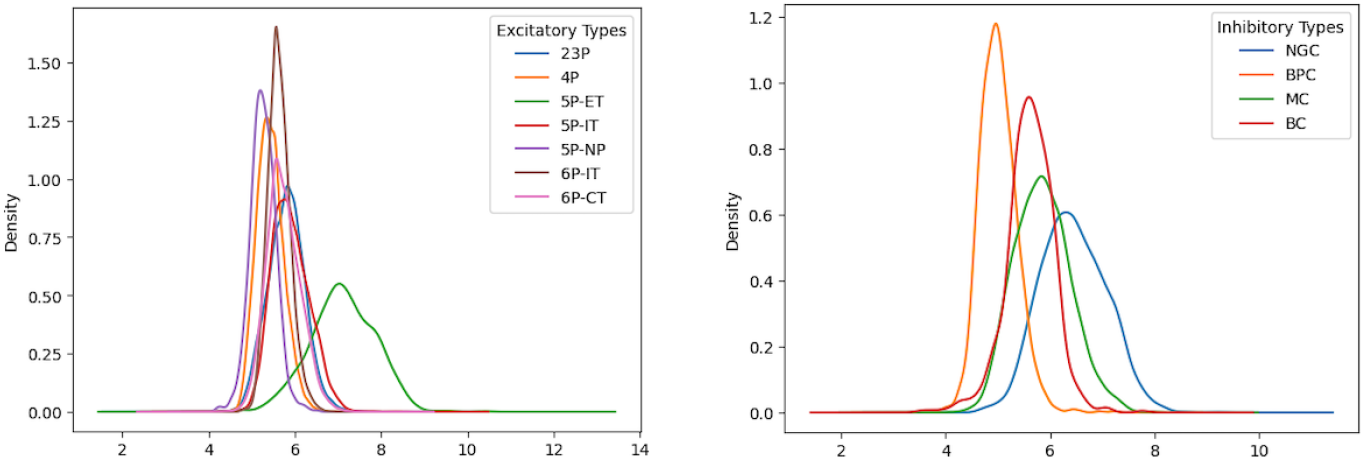
Distribution of soma radius. Estimated soma radius for Glutamatergic (left) and GABAergic (right) subclasses in the MICrONS dataset. The radius units (x-axis) are in *µ*m.

**Figure S12.**
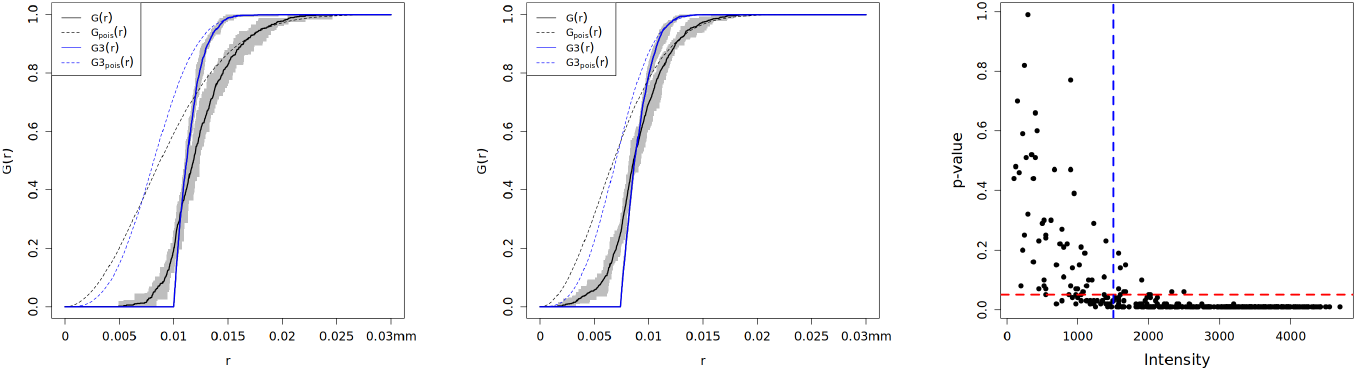
Simulation of 3D soma locations under spatial randomness. Nearest neighbor functions for the 3D and projected point patterns for average (left) and minimum (middle) soma radii used as hardcore distances. (Right) Decrease in statistical significance with respect to intensity of the projected point patterns. All units are in mm.

**Figure S13.**
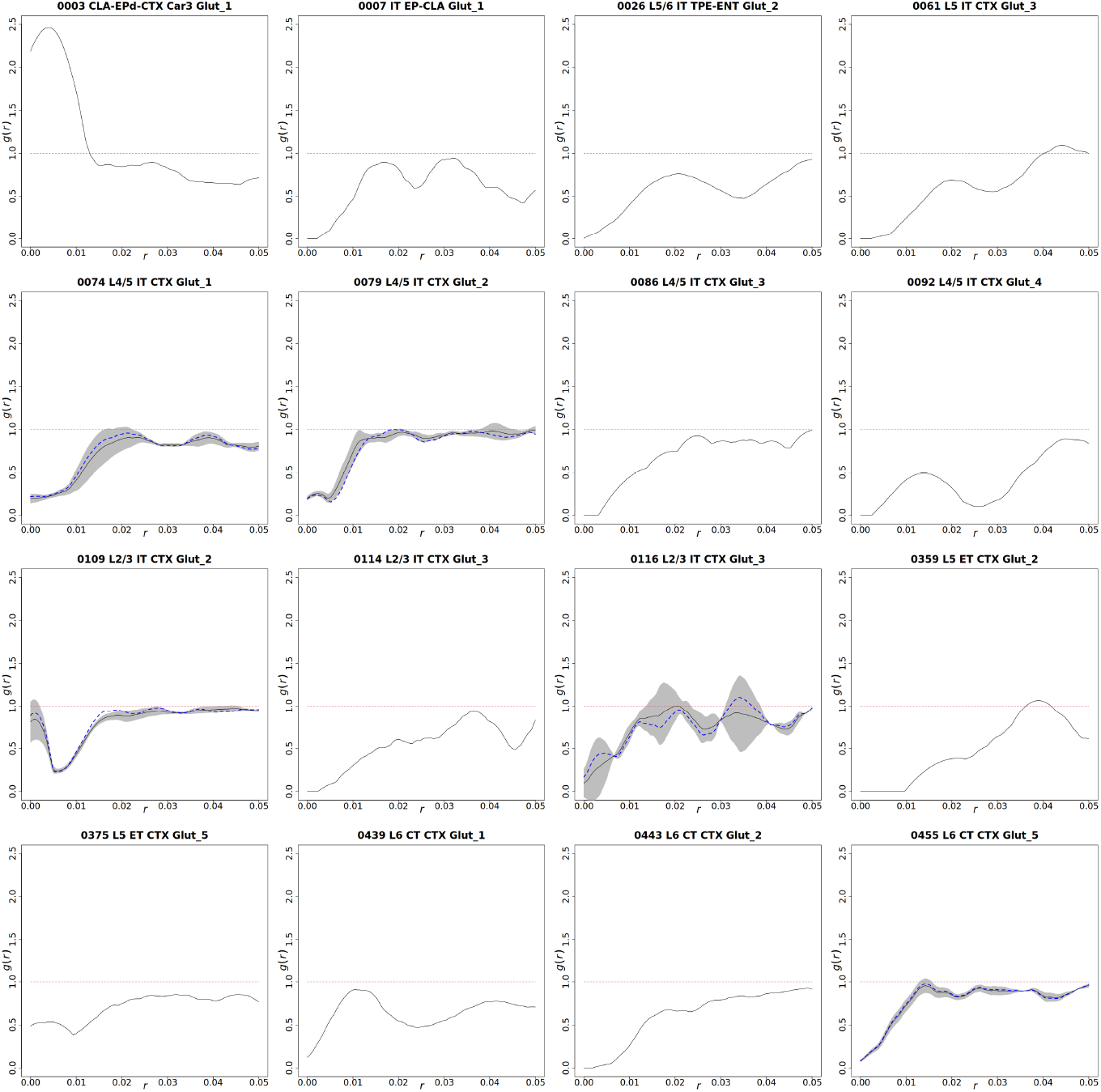
Pairwise correlation functions (PCF) for Glutamatergic clusters pooled over their corresponding sections that reject CSR. Solid lines are averaged PCFs of observed patterns plotted over the range of 50*µ*m, maximum value of `‘R’` for all excitatory clusters. The shaded regions cover lower and upper values over multiple sections. For cell types with multiple qualified sections, blue dotted lines corresponds to the section with lowest p-value for the type.

**Figure S14.**
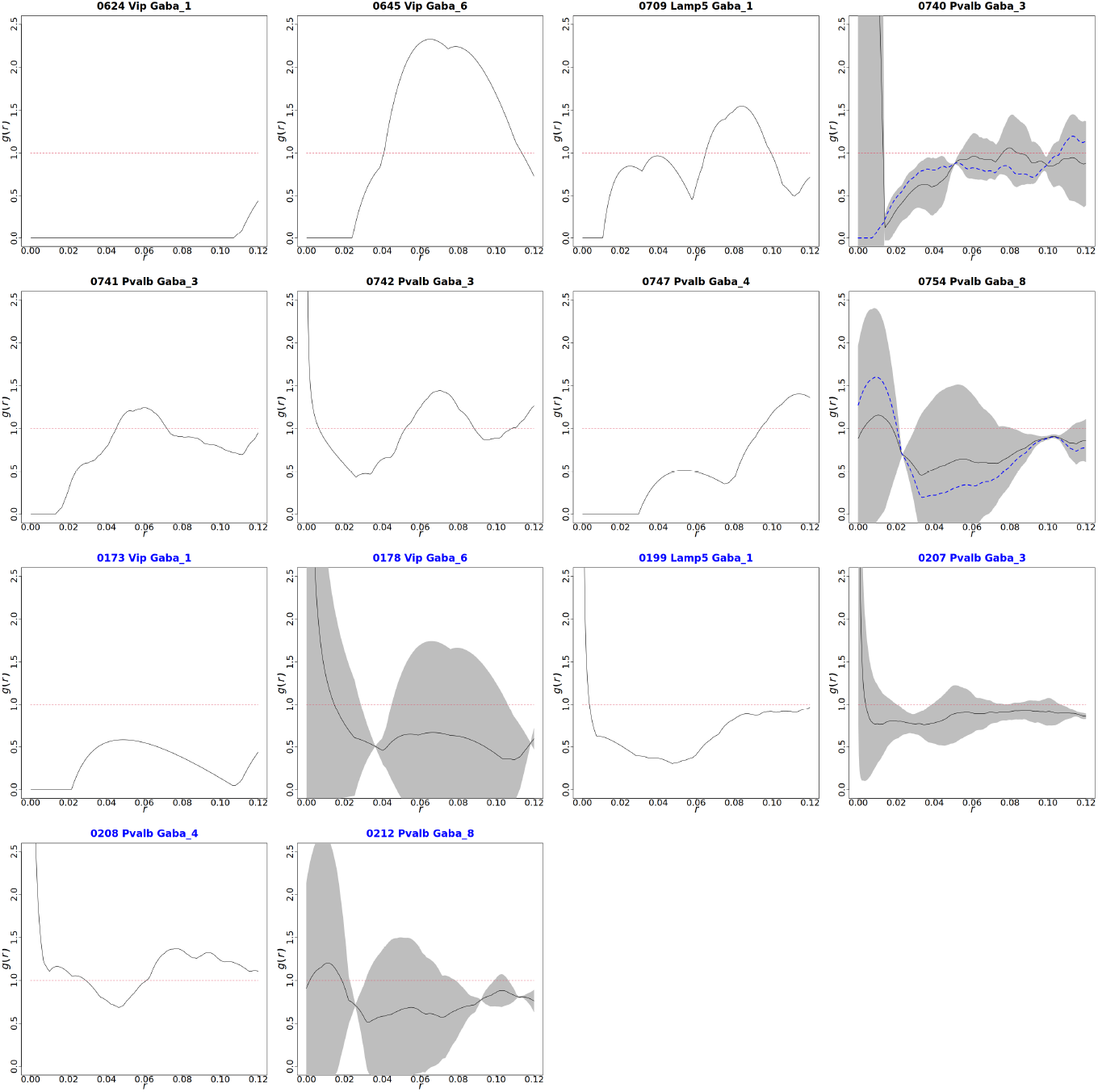
Pairwise correlation functions (PCF) for GABAergic clusters (black titles) and supertypes (blue titles) pooled over their corresponding sections that reject CSR. Solid lines are averaged PCFs of observed patterns plotted over the range of 120*µ*m, maximum value of `‘R’` for all inhibitory clusters. The shaded regions cover lower and upper values over multiple sections. For cell types with multiple qualified sections, blue dotted lines corresponds to the section with lowest p-value for the type.

**Figure S15.**
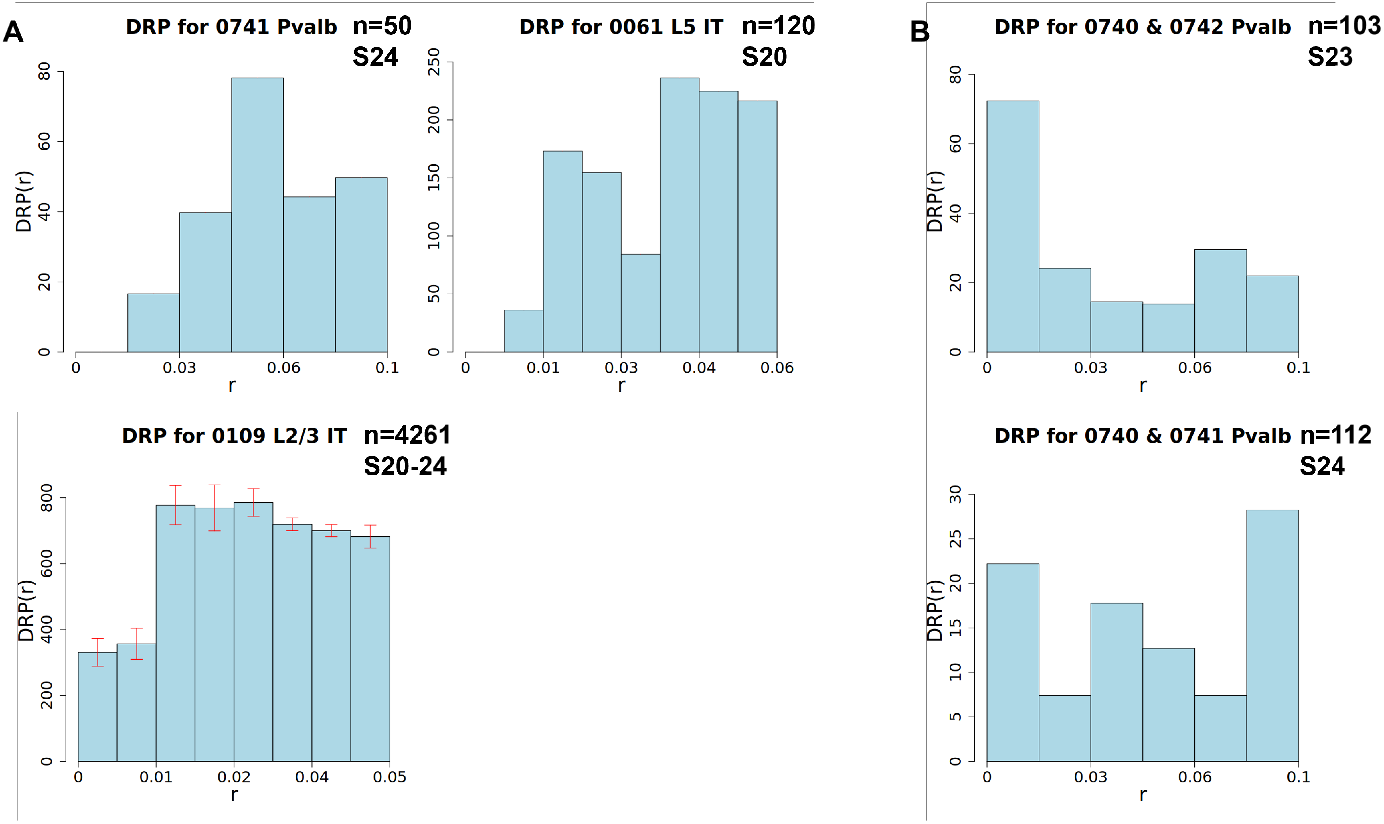
Density recovery profiles (DRPs) for selected individual types (A) and mixture populations (B). Compare, for instance, with Figure 2 in Ref.,^3^ Figure 1c in Ref.,^21^ or Figure 1E in Ref.^35^ Mixtures of inhibitory types display a clustering behavior at near distances, as previously reported at the subclass level.^17, 18^ The height of each bin represents the mean of the population. The error bars correspond to the standard deviation in each bin across sections (if the population in question passes the filtering steps in more than one sections). All units are in mm.

**Figure S16.**
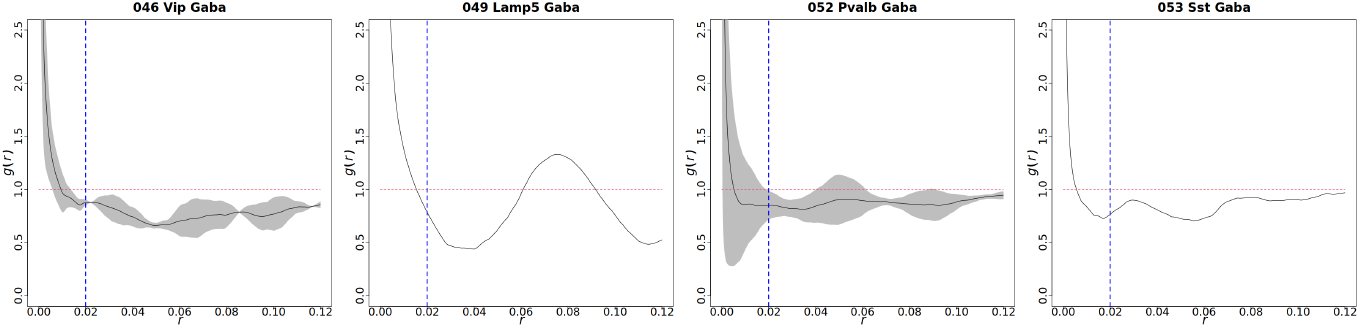
Pairwise correlation functions (PCF) for GABAergic subclasses pooled over their corresponding sections of interest. For details, see Figure S14. All subclasses show clustering effect for distances below approximately 20*µm* and spatial randomness for larger values of *r*.

**Figure S17.**
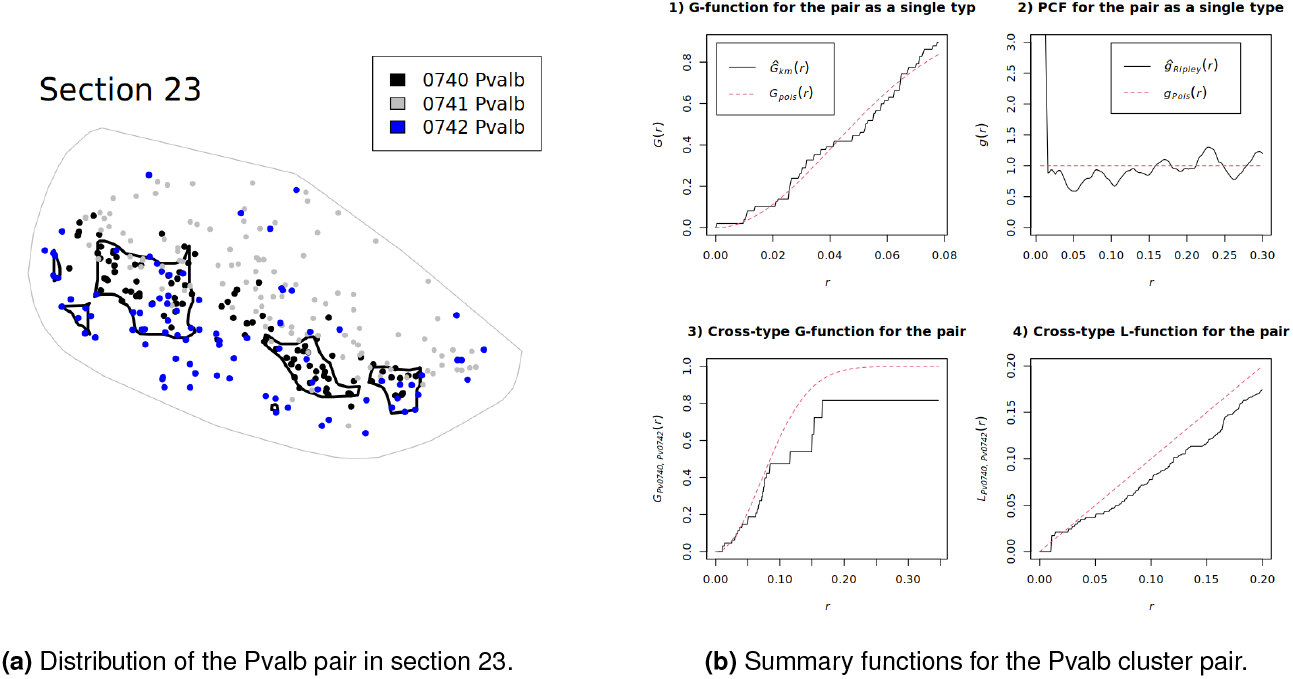
Case study for Pvalb cluster pair 0740 and 0742

**Figure S18.**
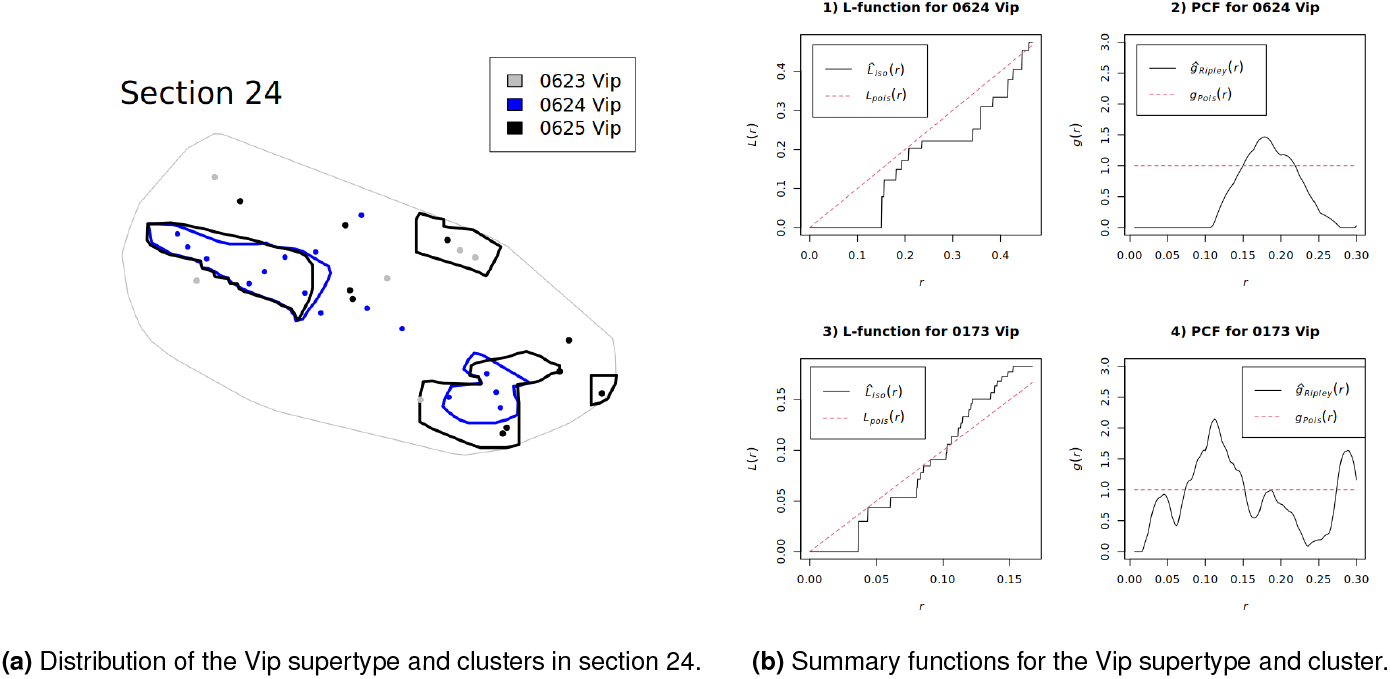
Case study for Vip supertype 0173 and cluster 0624.

**Figure S19.**
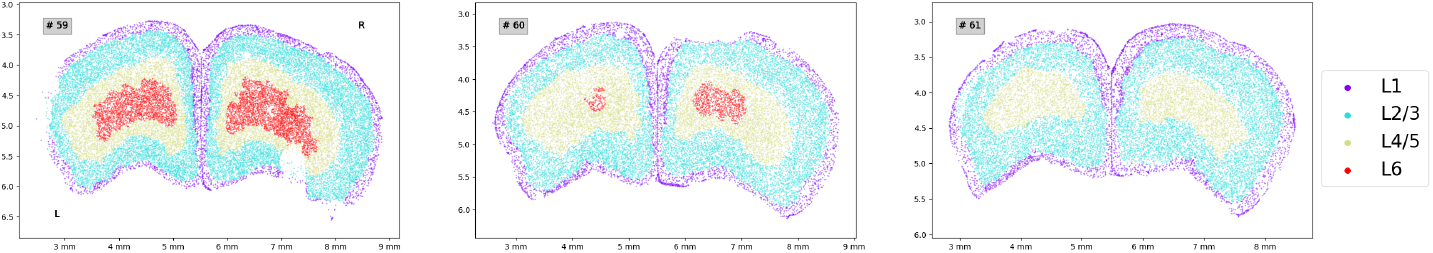
Annotated, detected cells in coronal sections at the dorsal part of the mouse brain used in this study, obtained from (24).

## Supplementary Tables

**Table S1.** Number of types after each filtering steps. Counts of cell types after each filtering steps for Glutamatergic (left) and GABAergic (right) types across 5 sections. Each column corresponds to one filtering step as follows. #label: label noise filter, #intensity: minimal intensity filter, #final: final counts after the secondary label noise filter.

| Section | #total | #label | #intensity | #final | Section | #total | #label | #intensity | #final |
| --- | --- | --- | --- | --- | --- | --- | --- | --- | --- |
| 20 | 159 | 17 | 9 | 8 | 20 | 124 | 21 | 5 | 4 |
| 21 | 148 | 14 | 10 | 9 | 21 | 112 | 19 | 3 | 3 |
| 22 | 150 | 14 | 4 | 2 | 22 | 105 | 14 | 2 | 2 |
| 23 | 112 | 18 | 10 | 9 | 23 | 103 | 12 | 2 | 2 |
| 24 | 79 | 15 | 8 | 7 | 24 | 83 | 11 | 5 | 5 |
| Total | 183 | 32 | 21 | 18 | Total | 133 | 28 | 11 | 10 |

**Table S2.** Filtering results for Glutamatergic and GABAergic clusters. The table shows the HC ratio (ratio of the number of high confidence cells to the total number of cells) per section from S20 (Section 20) to S24 (Section 24) for cell types passing the minimum intensity filter. Low HC ratio implies higher label noises. We highlight sections with HC ratio *≤* 0.5. In total, 18 Glutamatergic cell types and 10 GABAergic cell types are qualified in at least one section.

| Type | S20 | S21 | S22 | S23 | S24 |
| --- | --- | --- | --- | --- | --- |
| 1 0003 CLA-EPd-CTX Car3 Glut_1 | 0.64 | 0.68 |  |  |  |
| 2 0004 CLA-EPd-CTX Car3 Glut_1 |  | 0.79 |  | 0.74 |  |
| 3 0007 IT EP-CLA Glut_1 | 0.92 |  |  |  |  |
| 4 0026 L5/6 IT TPE-ENT Glut_2 |  |  |  | 0.55 |  |
| 5 0061 L5 IT CTX Glut_3 | 0.73 |  |  |  |  |
| 6 0067 L4/5 IT CTX Glut_1 |  |  |  | **0.47** |  |
| 7 0074 L4/5 IT CTX Glut_1 | 0.51 | 0.56 | 0.55 | 0.54 | 0.57 |
| 8 0079 L4/5 IT CTX Glut_2 | 0.62 | 0.58 | **0.44** |  | 0.51 |
| 9 0080 L4/5 IT CTX Glut_2 |  |  |  |  | **0.40** |
| 10 0086 L4/5 IT CTX Glut_3 |  |  |  | 0.92 | 0.89 |
| 11 0092 L4/5 IT CTX Glut_4 |  |  |  | 0.51 | 0.53 |
| 12 0109 L2/3 IT CTX Glut_2 | 0.85 | 0.86 | 0.81 | 0.82 | 0.80 |
| 13 0114 L2/3 IT CTX Glut_3 |  |  |  |  | 0.69 |
| 14 0116 L2/3 IT CTX Glut_3 |  | 0.51 |  | 0.71 | 0.72 |
| 15 0359 L5 ET CTX Glut_2 |  |  |  | 0.62 |  |
| 16 0375 L5 ET CTX Glut_5 |  |  |  | 0.69 |  |
| 17 0436 L6 CT CTX Glut_1 |  | **0.33** | **0.32** |  |  |
| 18 0439 L6 CT CTX Glut_1 | **0.50** | 0.59 |  |  |  |
| 19 0443 L6 CT CTX Glut_2 |  | 0.60 |  |  |  |
| 20 0453 L6 CT CTX Glut_5 | 0.76 |  |  |  |  |
| 21 0455 L6 CT CTX Glut_5 | 0.57 | 0.66 |  |  |  |
| 1 0624 Vip Gaba_1 |  |  |  |  | 0.75 |
| 2 0641 Vip Gaba_5 |  |  | 0.71 |  |  |
| 3 0645 Vip Gaba_6 | 0.83 |  |  |  | 0.74 |
| 4 0709 Lamp5 Gaba_1 |  |  | 0.73 |  |  |
| 5 0740 Pvalb Gaba_3 | 0.69 | 0.62 |  | 0.80 | 0.59 |
| 6 0741 Pvalb Gaba_3 |  |  |  |  | 0.71 |
| 7 0742 Pvalb Gaba_3 |  |  |  | 0.85 | 0.73 |
| 8 0743 Pvalb Gaba_4 | **0.48** |  |  |  |  |
| 9 0747 Pvalb Gaba_4 | 0.76 |  |  |  |  |
| 10 0754 Pvalb Gaba_8 | 0.86 | 0.74 |  |  |  |
| 11 0811 Sst Gaba_13 |  | 0.65 |  |  |  |

**Table S3.**
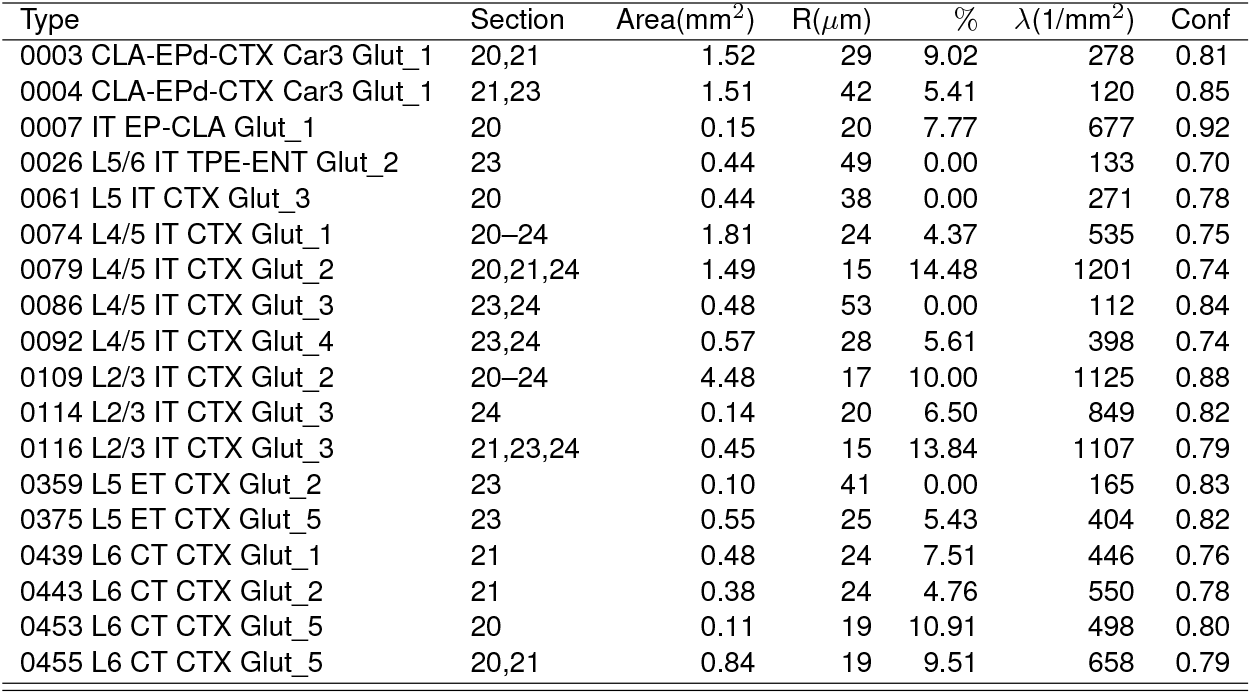
Statistics of Glutamatergic clusters before imposing an upper limit on the intensity of the selected region. For details, see Table 1.

| Type | Section | Area(mm<sup>2</sup>) | R(μm) | % | $\lambda$(1/mm<sup>2</sup>) | Conf |
| --- | --- | --- | --- | --- | --- | --- |
| 0003 CLA-EPd-CTX Car3 Glut_1 | 20,21 | 1.52 | 29 | 9.02 | 278 | 0.81 |
| 0004 CLA-EPd-CTX Car3 Glut_1 | 21,23 | 1.51 | 42 | 5.41 | 120 | 0.85 |
| 0007 IT EP-CLA Glut_1 | 20 | 0.15 | 20 | 7.77 | 677 | 0.92 |
| 0026 L5/6 IT TPE-ENT Glut_2 | 23 | 0.44 | 49 | 0.00 | 133 | 0.70 |
| 0061 L5 IT CTX Glut_3 | 20 | 0.44 | 38 | 0.00 | 271 | 0.78 |
| 0074 L4/5 IT CTX Glut_1 | 20–24 | 1.81 | 24 | 4.37 | 535 | 0.75 |
| 0079 L4/5 IT CTX Glut_2 | 20,21,24 | 1.49 | 15 | 14.48 | 1201 | 0.74 |
| 0086 L4/5 IT CTX Glut_3 | 23,24 | 0.48 | 53 | 0.00 | 112 | 0.84 |
| 0092 L4/5 IT CTX Glut_4 | 23,24 | 0.57 | 28 | 5.61 | 398 | 0.74 |
| 0109 L2/3 IT CTX Glut_2 | 20–24 | 4.48 | 17 | 10.00 | 1125 | 0.88 |
| 0114 L2/3 IT CTX Glut_3 | 24 | 0.14 | 20 | 6.50 | 849 | 0.82 |
| 0116 L2/3 IT CTX Glut_3 | 21,23,24 | 0.45 | 15 | 13.84 | 1107 | 0.79 |
| 0359 L5 ET CTX Glut_2 | 23 | 0.10 | 41 | 0.00 | 165 | 0.83 |
| 0375 L5 ET CTX Glut_5 | 23 | 0.55 | 25 | 5.43 | 404 | 0.82 |
| 0439 L6 CT CTX Glut_1 | 21 | 0.48 | 24 | 7.51 | 446 | 0.76 |
| 0443 L6 CT CTX Glut_2 | 21 | 0.38 | 24 | 4.76 | 550 | 0.78 |
| 0453 L6 CT CTX Glut_5 | 20 | 0.11 | 19 | 10.91 | 498 | 0.80 |
| 0455 L6 CT CTX Glut_5 | 20,21 | 0.84 | 19 | 9.51 | 658 | 0.79 |

**Table S4.** p-values for Glutamatergic individual type-section combinations at cell type level. `‘pv.adj’` denotes the Banjamini-Hochberg corrected p-values.

| Type | S24 | S23 | S22 | S21 | S20 | p-value | pv.adj |
| --- | --- | --- | --- | --- | --- | --- | --- |
| 0109 L2/3 IT CTX Glut_2 | 0.005 | 0.005 | 0.005 | 0.005 | 0.005 | 7.5e-08 | 1.4e-06 |
| 0074 L4/5 IT CTX Glut_1 | 0.005 | 0.005 | 0.005 | 0.072 | 0.008 | 1.1e-06 | 9.9e-06 |
| 0079 L4/5 IT CTX Glut_2 | 0.013 |  |  | 0.005 | 0.005 | 4.1e-05 | 2.5e-04 |
| 0455 L6 CT CTX Glut_5 |  |  |  | 0.005 | 0.007 | 3.9e-04 | 1.8e-03 |
| 0116 L2/3 IT CTX Glut_3 | 0.007 | 0.006 |  | 0.337 |  | 1.1e-03 | 4.0e-03 |
| 0092 L4/5 IT CTX Glut_4 | 0.005 | 0.067 |  |  |  | 3.0e-03 | 9.0e-03 |
| 0061 L5 IT CTX Glut_3 |  |  |  |  | 0.005 | 5.0e-03 | 0.01 |
| 0114 L2/3 IT CTX Glut_3 | 0.005 |  |  |  |  | 5.0e-03 | 0.01 |
| 0443 L6 CT CTX Glut_2 |  |  |  | 0.005 |  | 5.0e-03 | 0.01 |
| 0086 L4/5 IT CTX Glut_3 | 0.058 | 0.014 |  |  |  | 6.6e-03 | 0.012 |
| 0375 L5 ET CTX Glut_5 |  | 0.011 |  |  |  | 0.011 | 0.018 |
| 0026 L5/6 IT TPE-ENT Glut_2 |  | 0.015 |  |  |  | 0.015 | 0.022 |
| 0439 L6 CT CTX Glut_1 |  |  |  | 0.024 |  | 0.024 | 0.033 |
| 0007 IT EP-CLA Glut_1 |  |  |  |  | 0.027 | 0.027 | 0.035 |
| 0359 L5 ET CTX Glut_2 |  | 0.039 |  |  |  | 0.039 | 0.047 |
| 0003 CLA-EPd-CTX Car3 Glut_1 |  |  |  | 0.922 | 0.008 | 0.044 | 0.049 |
| 0453 L6 CT CTX Glut_5 |  |  |  |  | 0.061 | 0.061 | 0.065 |
| 0004 CLA-EPd-CTX Car3 Glut_1 |  | 0.064 |  | 0.481 |  | 0.14 | 0.14 |

**Table S5.** p-values for individual GABAergic type-section combinations at the cell type and supertype levels. `‘pv.adj’` denotes the Banjamini-Hochberg corrected p-values.

| Type/Supertype | S24 | S23 | S22 | S21 | S20 | p-value | pv.adj |
| --- | --- | --- | --- | --- | --- | --- | --- |
| 0754 Pvalb Gaba_8 |  |  |  | 0.008 | 0.028 | 0.0021 | 0.02 |
| 0212 Pvalb Gaba_8 |  |  |  | 0.010 | 0.026 | 0.0024 | 0.02 |
| 0624 Vip Gaba_1 | 0.011 |  |  |  |  | 0.011 | 0.024 |
| 0641 Vip Gaba_5 |  |  | 0.013 |  |  | 0.013 | 0.024 |
| 0645 Vip Gaba_6 | 0.025 |  |  |  | 0.085 | 0.015 | 0.024 |
| 0740 Pvalb Gaba_3 | 0.026 | 0.039 |  | 0.841 | 0.017 | 0.0044 | 0.024 |
| 0741 Pvalb Gaba_3 | 0.007 |  |  |  |  | 0.007 | 0.024 |
| 0742 Pvalb Gaba_3 | 0.063 | 0.040 |  |  |  | 0.018 | 0.024 |
| 0747 Pvalb Gaba_4 |  |  |  |  | 0.017 | 0.017 | 0.024 |
| 0811 Sst Gaba_13 |  |  |  | 0.018 |  | 0.018 | 0.024 |
| 0173 Vip Gaba_1 | 0.013 |  |  |  |  | 0.013 | 0.024 |
| 0178 Vip Gaba_6 | 0.020 |  |  |  | 0.091 | 0.013 | 0.024 |
| 0226 Sst Gaba_13 |  |  |  | 0.018 |  | 0.018 | 0.024 |
| 0709 Lamp5 Gaba_1 |  |  | 0.029 |  |  | 0.029 | 0.035 |
| 0199 Lamp5 Gaba_1 |  |  | 0.068 |  |  | 0.068 | 0.077 |
| 0208 Pvalb Gaba_4 |  |  |  |  | 0.101 | 0.100 | 0.11 |
| 0207 Pvalb Gaba_3 | 0.017 | 0.994 |  | 0.930 | 0.126 | 0.13 | 0.13 |

**Table S6.**
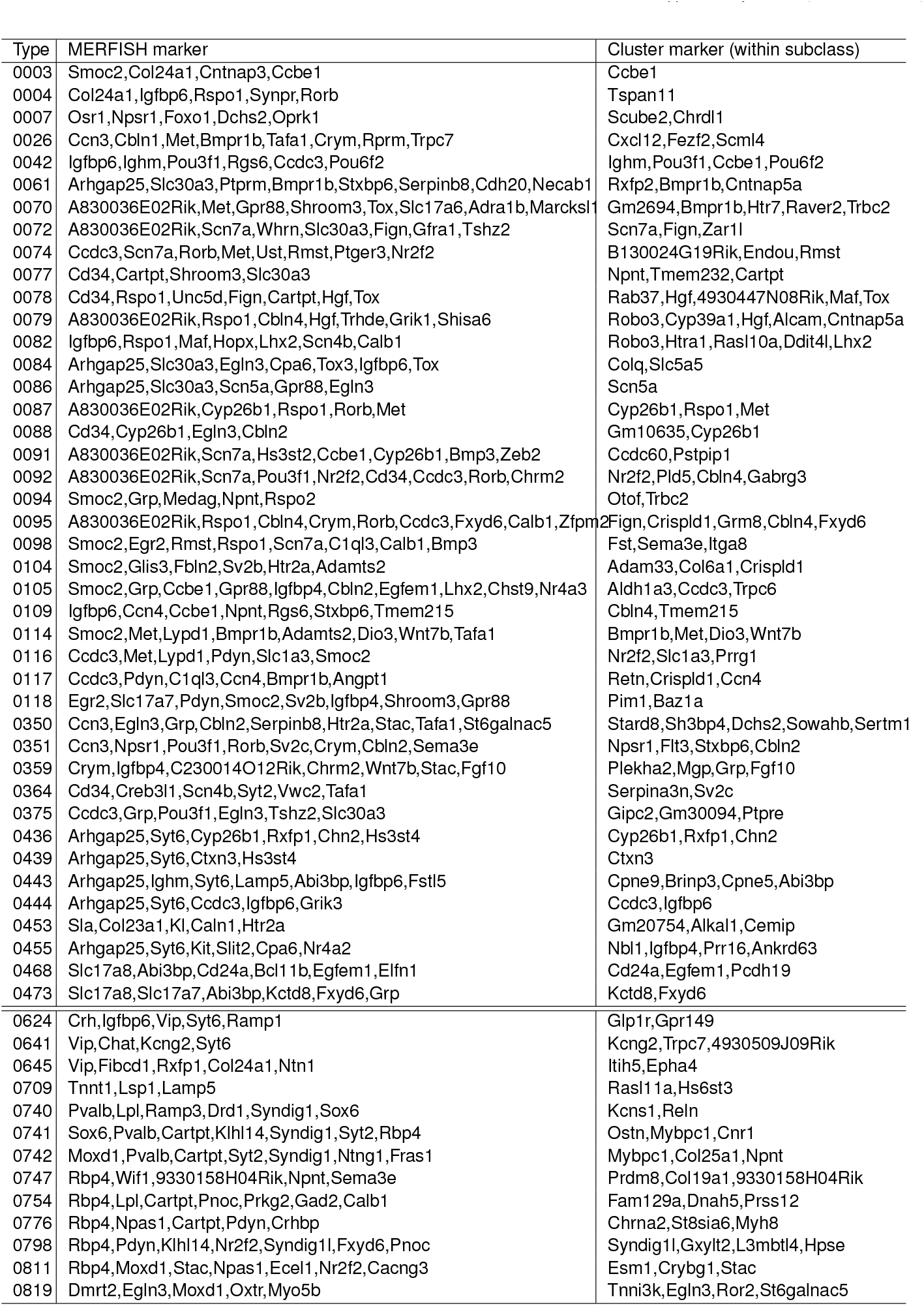
List of differentially expressed genes in studied cell types. For readability, cell type names are abbreviated for type number only. MERFISH marker: Unique combination of markers present on MERFISH gene panel seperating from all the other clusters. Cluster marker (within subclass): Unique combination of markers separating from other clusters within the same subclass. Compiled from the full table at https://alleninstitute.github.io/abc_atlas_access/descriptions/WMB-taxonomy.html.

**Table S7.** All GABAergic subclasses display clustering effects. p-values across different brain sections are combined by Fisher’s method and corrected for multiple testing using the Benjamini-Hochberg method. The first p-value column corresponds to the one-sided DCLF test for clustering. The second p-value column corresponds to the one-sided DCLF test for repulsion.

| Subclass | Section | p-value1 | p-value2 | Area(mm<sup>2</sup>) | R(μm) | % | $\lambda$(1/mm<sup>2</sup>) | Conf |
| --- | --- | --- | --- | --- | --- | --- | --- | --- |
| 046 Vip Gaba | 20,24 | 7.9e-03 | 0.12 | 4.81 | 46 | 0.76 | 42 | 0.76 |
| 049 Lamp5 Gaba | 22 | 0.005 | 0.63 | 2.70 | 59 | 2.47 | 30 | 0.77 |
| 052 Pvalb Gaba | 20,21,23,24 | 1.1e-06 | 0.51 | 12.32 | 30 | 3.41 | 93 | 0.78 |
| 053 Sst Gaba | 21 | 0.005 | 0.24 | 3.08 | 39 | 2.14 | 61 | 0.73 |

**Table S8.** p-values for individual GABAergic cluster-section combinations at the subclass level.

| Subclass name | S24 | S23 | S22 | S21 | S20 | p-value |
| --- | --- | --- | --- | --- | --- | --- |
| 046 Vip Gaba | 0.16 |  |  |  | 0.159 | 0.12 |
| 049 Lamp5 Gaba |  |  | 0.63 |  |  | 0.63 |
| 052 Pvalb Gaba | 0.085 | 0.998 |  | 1 | 0.318 | 0.51 |
| 053 Sst Gaba |  |  |  | 0.24 |  | 0.24 |

**Table S9.** Table of cell counts by parcellation structures for mixture of Pvalb 0740 and Pvalb 0742 in Section 23.

| Parcellation | Total | Pvalb 0740 | Pvalb 0742 |
| --- | --- | --- | --- |
| AUDd2/3 | 5 | 0 | 5 |
| AUDd4 | 1 | 0 | 1 |
| AUDp4 | 11 | 8 | 3 |
| AUDp5 | 30 | 26 | 4 |
| AUDpo2/3 | 6 | 2 | 4 |
| AUDpo4 | 5 | 3 | 2 |
| SSs2/3 | 30 | 3 | 27 |
| SSs4 | 34 | 23 | 11 |
| SSs5 | 9 | 8 | 1 |
| TEa4 | 4 | 2 | 2 |
| VISC4 | 5 | 5 | 0 |
| VISC5 | 12 | 6 | 6 |
| VISpor2/3 | 3 | 0 | 3 |

**Table S10.** A majority of Glutamatergic clusters and all GABAergic clusters display homotypic avoidance. The table summarizes the second dataset used in this study, which covers the coronal sections at the dorsal part of the mouse brain, obtained from (24). See Table 1 for notation details.

| Cell type | Section | p-value | Area(mm<sup>2</sup>) | R(μm) | % | $\lambda$(1/mm<sup>2</sup>) | conf | n |
| --- | --- | --- | --- | --- | --- | --- | --- | --- |
| 0072 L4/5 IT CTX Glut_1 | 59–61 | 9.0e-05 | 1.25 | 30 | 0.66 | 447 | 0.72 | 558 |
| 0084 L4/5 IT CTX Glut_3 | 59–61 | 9.0e-05 | 1.28 | 38 | 0.80 | 335 | 0.71 | 430 |
| 0091 L4/5 IT CTX Glut_4 | 59–61 | 9.0e-05 | 0.97 | 31 | 1.31 | 534 | 0.71 | 521 |
| 0095 L4/5 IT CTX Glut_5 | 59–61 | 9.0e-05 | 1.69 | 29 | 0.78 | 426 | 0.72 | 751 |
| 0105 L2/3 IT CTX Glut_1 | 59–61 | 9.0e-05 | 0.25 | 19 | 1.29 | 1216 | 0.74 | 298 |
| 0104 L2/3 IT CTX Glut_1 | 59–61 | 9.6e-05 | 0.35 | 16 | 3.83 | 875 | 0.72 | 324 |
| 0117 L2/3 IT CTX Glut_4 | 59–61 | 9.6e-05 | 0.62 | 32 | 2.07 | 560 | 0.71 | 323 |
| 0082 L4/5 IT CTX Glut_2 | 59–61 | 1.1e-04 | 0.54 | 38 | 0.00 | 423 | 0.71 | 236 |
| 0351 L5 ET CTX Glut_1 | 59–61 | 1.1e-04 | 1.53 | 36 | 0.34 | 282 | 0.72 | 431 |
| 0078 L4/5 IT CTX Glut_2 | 59–61 | 3.0e-04 | 0.76 | 34 | 1.42 | 279 | 0.73 | 210 |
| 0077 L4/5 IT CTX Glut_2 | 60, 61 | 5.6e-04 | 0.45 | 32 | 1.39 | 427 | 0.70 | 194 |
| 0087 L4/5 IT CTX Glut_3 | 60, 61 | 5.6e-04 | 0.74 | 34 | 1.32 | 317 | 0.72 | 243 |
| 0436 L6 CT CTX Glut_1 | 59, 60 | 5.6e-04 | 0.84 | 19 | 1.38 | 737 | 0.74 | 630 |
| 0098 L4/5 IT CTX Glut_5 | 59, 60 | 1.1e-03 | 0.48 | 30 | 1.45 | 369 | 0.70 | 175 |
| 0042 L6 IT CTX Glut_2 | 59, 60 | 1.6e-03 | 0.76 | 29 | 0.98 | 318 | 0.70 | 260 |
| 0070 L4/5 IT CTX Glut_1 | 59 | 7.4e-03 | 0.30 | 44 | 2.20 | 302 | 0.70 | 91 |
| 0444 L6 CT CTX Glut_2 | 59 | 7.4e-03 | 0.46 | 35 | 0.00 | 329 | 0.71 | 150 |
| 0094 L4/5 IT CTX Glut_4 | 61 | 9.7e-03 | 0.30 | 19 | 0.00 | 359 | 0.70 | 107 |
| 0118 L2/3 IT CTX Glut_4 | 59 | 0.013 | 0.31 | 26 | 0.00 | 338 | 0.69 | 105 |
| 0350 L5 ET CTX Glut_1 | 59 | 0.025 | 0.11 | 29 | 0.00 | 367 | 0.77 | 41 |
| 0061 L5 IT CTX Glut_3 | 59 | 0.042 | 0.38 | 33 | 3.96 | 268 | 0.68 | 101 |
| 0364 L5 ET CTX Glut_2 | 59 | 0.055 | 0.55 | 46 | 8.93 | 205 | 0.70 | 112 |
| 0468 L5 NP CTX Glut_3 | 61 | 0.14 | 0.48 | 45 | 0.00 | 132 | 0.75 | 63 |
| 0088 L4/5 IT CTX Glut_3 | 59 | 0.26 | 0.40 | 43 | 5.48 | 182 | 0.71 | 73 |
| 0473 L5 NP CTX Glut_4 | 60 | 0.30 | 0.62 | 60 | 4.44 | 73 | 0.70 | 45 |
| 0740 Pvalb Gaba_3 | 61 | 0.012 | 0.92 | 65 | 0 | 123 | 0.71 | 113 |
| 0742 Pvalb Gaba_3 | 61 | 0.012 | 1.29 | 81 | 0 | 63 | 0.71 | 82 |
| 0776 Sst Gaba_4 | 60 | 0.017 | 0.73 | 120 | 0 | 18 | 0.71 | 13 |
| 0819 Sst Gaba_16 | 59 | 0.024 | 1.60 | 120 | 0 | 9 | 0.69 | 15 |
| 0798 Sst Gaba_9 | 59 | 0.041 | 1.32 | 120 | 0 | 13 | 0.70 | 17 |

## Notes

https://alleninstitute.github.io/abc_atlas_access/

### Competing Interest Statement

The authors have declared no competing interest.

### Summary of Updates

Added results on a second dataset. Supplemental material updated.

